# An RNA binding protein, RNP1A, works with Contractility Kit proteins to facilitate macropinocytosis

**DOI:** 10.1101/2022.07.08.499268

**Authors:** Yinan Liu, Jessica Leng, Ly TS Nguyen, Douglas N. Robinson

**Affiliations:** Departments of Cell Biology, Johns Hopkins School of Medicine, Baltimore, MD 21205; Departments of Pharmacology and Molecular Sciences, Johns Hopkins School of Medicine, Baltimore, MD 21205; Case Western Reserve University School of Medicine, Cleveland, Ohio, 44106

**Keywords:** Contractility Kits, mechanosensing, macropinocytosis, RNA binding proteins

## Abstract

Cell shape regulation is important for many biological processes. Some cell shape regulating proteins harbor mechanoresponsive properties that enable them to sense and respond to mechanical cues, allowing for cell adaptation. In *Dictyostelium discoideum*, mechanoresponsive network proteins include Cortexillin I and IQGAP1, which assemble in the cytoplasm into macromolecular complexes, which we term Contractility Kits. *In vivo* fluorescence cross-correlation spectroscopy revealed that Cortexillin I also interacts with an RNA-binding protein, RNP1A. The *rnp1A* knockdown cells have reduced cell growth rate, reduced adhesion, defective cytokinesis, and a gene expression profile that indicates *rnp1A* knockdown cells shift away from the vegetative growth state. RNP1A binds to transcripts encoding proteins involved in macropinocytosis. One of these, DlpA, facilitates macropinosome maturation, similar to RNP1A. Loss of different CK proteins leads to macropinocytotic defects characterized by reduced macropinocytotic crown size. RNP1A interacts with IQGAP1 *in vivo* and has cross-talk with IQGAP1 during macropinocytosis. Overall, RNP1A contributes to macropinocytosis, in part through interacting with transcripts encoding macropinocytotic proteins like *dlpA*, and does so in coordination with the Contractility Kit proteins.

## Introduction

Cell shape control is critical to many biological processes, including proliferation, differentiation, cell migration, endocytosis, and exocytosis (Clark & Paluch, 2011; Kaverina & Straube, 2011; Sapala, Runions et al., 2019; Srivastava, Iglesias et al., 2016; Dai & Sheetz, 1995; Joseph & Liu, 2020). The molecular basis of cell shape regulation is dependent on several cellular systems, including cytoskeletal polymers, motor proteins, adhesion proteins, signaling proteins and membrane channels. These systems collaborate to regulate and actively drive cell shape change, as well as respond to external stimuli allowing for cellular response and adaptation (Kaverina & Straube, 2011; McBeath, Pirone et al., 2004; Morlot, Galli et al., 2012 Rape, Guo et al., 2011; Pollard & Cooper, 2009; Jiang & Sun, 2013). External stimuli causing mechanical stress or mechanical cues, are received by this molecular infrastructure and influence cellular output. In particular, some proteins sense and respond to external mechanical stimuli, by translocating, modulating local protein architecture, and triggering downstream signaling pathways (Kasza & Zallen, 2011; Ondeck, Kumar et al., 2019; McWhorter, Wang et al., 2013; Luo, Mohan et al., 2013; Schiffhauer, Luo et al., 2016). These proteins are coined as mechanoresponsive proteins because they sense and redistribute to sites where mechanical stress has been imposed. In *Dictyostelium discoideum*, several mechanoresponsive proteins have been identified, including the motor protein Myosin II, the actin cross-linker Cortexillin I, and the regulatory protein IQGAP2 (Kee, Ren et al., 2012; Luo et al., 2013; Ren, Effler et al., 2009). More recently, it has been discovered that these mechanoresponsive proteins assemble into macromolecular complexes, which we term Contractility Kits (CKs), in the cytoplasm before mechanosensation. We have also discovered that CKs containing IQGAP2 define mechanoresponsive CKs, but a sister protein IQGAP1 serves as a negative regulator such that the CKs are non-mechanoresponsive without IQGAP2. The organization of these proteins into the CKs likely then helps facilitate robust and rapid responses to external mechanical stimuli imposed upon the cell cortex (Kothari, Srivastava et al., 2019).

Previously, we genetically identified an RNA-binding protein, RNP1A (DDB ID:DDB_G0284167), as a suppressor of nocodazole (Zhou, Kee et al., 2010; Ngo, Miao et al., 2016). Through follow up studies, we found RNP1A interacts with Cortexillin I (Kothari et al., 2019). *In vivo* affinity measurements using Fluorescence Cross-Correlation Spectroscopy (FCCS) indicates that RNP1A is one of the strongest biochemical interactors with Cortexillin I, having an apparent *in vivo* K_D_ of 0.33 μM (Kothari et al., 2019). Coincidentally, in the same study, another RNA-binding protein (RNP1B; DDB ID:DDB_G0285361) was also identified as an interactor of Cortexillin I through a proteomics study. Collectively, these findings raised the question about the functional relationship between RNA-binding proteins and CK proteins.

RNA-binding proteins (RBPs) have a myriad of functions in cells, including regulating RNA export, modification, localization, stability, and translation, as well as regulating transcription and chromatin remodeling. Many RBPs have multiple functions, which are dependent on its binding partners, imposed cellular stresses, and the biological processes in which the RBP is acting (Kishore, Luber et al., 2010; Balcerak, Trebinska-Stryjewska et al., 2019; Kilchert, Strasser et al., 2020). In *Dictyostelium discoideum*, emerging evidence has linked RBP functions to cell shape regulation. During chemotaxis, RBP Puf118 facilitates localization of RNA transcripts encoding proteins that are required for chemotaxis and that localize to the chemotaxing front (Hotz & Nelson, 2017). On the other hand, the *adenylyl cyclase A* RNA transcript localizes to the rear of chemotaxing cells and is locally translated to facilitate the adenylyl cyclase A protein’s role at the rear of a chemotaxing cell (Wang, Chen et al., 2018). RBPs have also been implicated in regulating cell shape and migration in other organisms. For example, RBPs facilitate sub-cellular localization and translation of β-actin mRNA in neuronal growth cones (Zhang, Eom et al., 2001), as well as localized mRNA translation at the cell migration front (Dermit, Dodel et al., 2020).

Here, we set out to identify the function of RNP1A in cell shape regulation and its functional interactions with the mechanoresponsive system and the Contractility Kit proteins. We found that *rnp1A* knockdown cells have slower cell growth, decreased cell adhesion, and cytokinetic defects. RNP1A localizes to the protrusive front during random cell migration and interacts with IQGAP1. RNP1A is slightly mechanoresponsive and contributes to maintaining cell cortical tension. Moreover, *rnp1A* knockdown leads to a major reduction in gene expression of translation and macropinocytosis pathways, which are critical to *Dictyostelium* vegetative growth. *Dictyostelium* normally undergoes a vegetative growth cycle; however, when nutrients become scarce, *Dictyostelium* enters a development stage (Flowers, Li et al., 2010). From gene expression profiling, genes enriched in *Dictyostelium* development stage are upregulated upon *rnp1A* knockdown. Analysis of transcripts bound by RNP1A revealed a large fraction (9 out of 24 of the RNAs identified) that encode proteins involved in macropinocytosis. Significantly, one such protein DlpA has also been found to maintain normal actomyosin assembly at the cleavage furrow during cytokinesis (Masud Rana, Tsujioka et al., 2013). We assessed macropinocytosis activity in the *rnp1A* knockdown cells and found that macropinocytosis is significantly impaired in these cells. Both *rnp1A* knock down and *dlpA* null cells showed delayed loss of internalized TRITC-Dextran. Furthermore, we found that IQGAP1, Cortexillin I, and Myosin II contribute to maintaining the size of macropinocytotic crowns. Overall, RNP1A interacts with Contractility Kit proteins and transcripts encoding *dlpA* to facilitate macropinocytosis. In so doing, RNP1A promotes *Dictyostelium* vegetative growth.

## Results

### RNP1A is important for cell growth, adhesion and cytokinesis

To study the function of RNP1A, we attempted to create an *rnp1A* knock-out or knockdown cell line. We first attempted to use CRISPR (Sekine, Kawata et al., 2018) to knock out *rnp1A* with two different guide RNAs. However, we could only acquire in-frame deletion of three nucleotides (37-39), which did not lead to knock-out of *rnp1A*. We then switched to using a long RNA hairpin approach to knock down *rnp1A*. We had to use a very stringent drug selection scheme to acquire cells that expressed *rnp1A* hairpin. Overall, combining observations from CRISPR and long RNA hairpin knockdown, *rnp1A* appears to be essential for cell survival.

Upon acquiring *rnp1A* knockdown cells, we first characterized the degree of knock down by quantifying RNA transcript and protein amount (**Fig. 1A; Appendix Fig.1**). The *rnp1A* knockdown cell line exhibited a 78% reduction (median) in *rnp1A* transcript and a 52% reduction (median) in RNP1A protein. We also acquired RNP1A-overexpressing (RNP1A-OE) cells by exogenously expressing RNP1A (**Fig. 1A; Appendix Fig.1**). These cells showed an increase in *rnp1A* transcript level by 439% (median) and an increase in RNP1A protein level by 69% (median).

**Figure 1.**
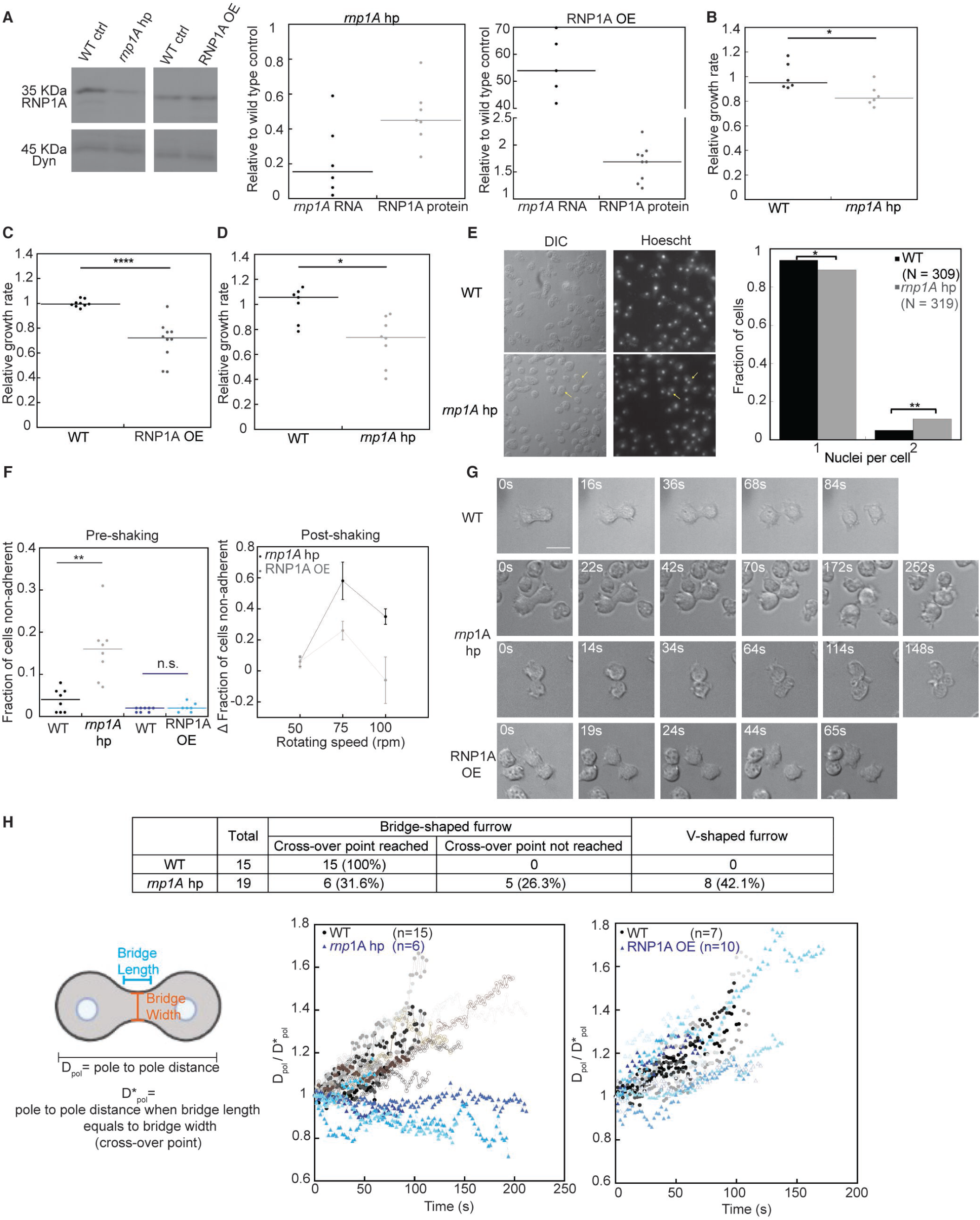
RNP1A is important for cell growth, adhesion and cytokinesis. (A) *rnp1A* knockdown and overexpressing levels were quantified by qRT-PCR and western blotting analysis. Dynacortin is a loading control. Western blot images were extracted from the same blot as shown in **Fig. 7A**. Original blots are shown in **Appendix Fig. 1**. Western blot data were analyzed by comparing to the total protein amount extracted from Coomassie staining from a replica gel. Each dot represents result from a single qRT-PCR run or a western blot. For *rnp1A* knockdown cells, qRT-PCR data were pooled from results from 4 different biological replicates and western blot data from 2 biological replicates. For RNP1A overexpressing cells, both qRT-PCR data and western blot data were pooled from results from 2 biological replicates. Growth rates quantified from (B) suspension culture of *rnp1A* knockdown cells, (C) suspension culture of overexpressing cells, and (D) *rnp1A* knockdown cells grown on substrate were normalized to that of WT control. Each dot represents relative growth rate quantified from a single flask or a single well. (E) Wild type control and *rnp1A* knockdown cells were fixed and nuclei were stained with Hoechst. Yellow arrows point to example cells containing two nuclei. Comparison of proportions test was performed to determine statistical difference. Data were pooled from 309 wild type control cells and *rnp1A* knockdown cells. Scale bar, 10 µm. (F) *rnp1A* knockdown cells and RNP1A overexpressing cells were shaken on rotating platform for 30 minutes. Pre-shaking fraction of non-adherent cells was calculated as cells suspended in medium over total number of cells. Each dot represents fraction calculated from a single well. Post-shaking fraction of non-adherent cells is presented as the incremental fraction of non-adherent cells compared to control. Error bars indicate standard errors. Data were pooled from three independent experiments. (G) DIC images of wild type control, *rnp1A* knockdown, or RNP1A overexpressing cells during the progression of cytokinesis. Scale bar, 10 µm. (H) Number of cells exhibiting either bridge-shaped or V-shaped cleavage furrow was counted for wild type control and *rnp1A* knockdown cells. Among cells that exhibited bridge-shaped cleavage furrow, number of cells were counted based on if they reached cross-over point (when bridge diameter equals bridge length). Pole-to-pole distances were quantified at and after cells reached cross-over point and normalized to pole-to-pole distance at cross-over point. For *rnp1A* knockdown cells, data were pooled from 15 wild type control cells and 6 *rnp1A* knockdown cells. For RNP1A overexpressing cells, data were pooled from 7 wild type control cells and 10 RNP1A overexpressing cells. Data information: All statistical analysis was done with Kruskal–Wallis followed by Wilcoxon-Mann–Whitney test, unless otherwise specified. *, P≤0.05; **, P≤0.01; ***, P≤0.001; ****, P≤0.0001; n.s, not significant.

We then characterized alterations in cell growth and found both *rnp1A* knockdown and overexpressing cell lines had decreased growth rates. We first grew cells in suspension culture and determined the relative cell growth rate for both cell lines (**Figs. 1B, 1C**). The *rnp1A* knockdown cell line had a growth rate of 0.83 (median) relative to the wild type parental control. The RNP1A-OE cell line’s growth rate was 0.72 (median) relative to control. In general, cell growth rate can be affected by a myriad of factors, including cell cycle progression, energy state, cytokinesis, and adhesion, which in turn can also impact cytokinesis fidelity (DeBerardinis, Lum et al., 2008; Kanada, Nagasaki et al., 2005).

From a cell mechanics regulation standpoint, we are especially interested in cytokinesis and adhesion ability. Interestingly, when we grew *rnp1A* knockdown cells on substrate, these cells exhibited even more severe cell growth defect (**Fig. 1D**). The relative growth rate for *rnp1A* knockdown cells on substrate was 0.74 (median) relative to control. This suggest that *rnp1A* knockdown cells could have a defect in adhesion ability. Moreover, when we measured the nuclei per cell count, which is an indicator of cytokinesis fidelity, we also observed an increase in the fraction of binucleated cells (wild type: 5%; *rnp1A* hairpin knockdown: 11%) (**Fig. 1E)**. The increase in fraction of binucleated cells implies that *rnp1A* hairpin knockdown cells have a mild cytokinetic defect.

We further explored changes in cell-substrate adhesion in *rnp1A* knockdown and RNP1A*-*OE cell lines. The *rnp1A* knockdown cells exhibited a higher fraction of non-adherent cells even before being subjected to shaking challenge, suggesting that *rnp1A* knockdown cells have a cell-substrate adhesion defect (**Fig. 1F**). After being subjected to shaking, a significant increase in the fraction of non-adherent cells was observed for *rnp1A* knockdown cells at 75 rpm and 100 rpm, whereas RNP1A-OE cells had a moderate increase at 75 rpm. In all, the adhesion assays indicate that *rnp1A* knockdown cells exhibit the most significant adhesion defect, and RNP1A-OE cells are only slightly defective in adhesion after shaking.

We then examined the process of cytokinesis in *rnp1A* knockdown and RNP1A-OE cells, to understand if the growth defects we observed could be attributed, in part, to a cytokinesis defect in these cell lines. We observed a significant cytokinetic abnormality in *rnp1A* knockdown cell line but not in RNP1A-OE cells (**Fig. 1G**). Overall, the cytokinesis morphology of *rnp1A* knockdown cells is significantly different than wild type cells (**Figs. 1G, 1H; Movies EV1, 2A, 2B**). First, all wild type cells assume a cylindrical bridge-shaped cleavage furrow, which represents the existence of the combination of ring constriction forces and traction forces (Jahan & Yumura, 2017). However, only 57.9% of *rnp1A* hairpin knockdown cells assumed a bridge-shaped furrow (wild type like). The remaining 42.1% of *rnp1A* knockdown cells assumed a V-shaped furrow, resembling cytokinesis A, which majorly depends on the constriction forces at the furrow and less on traction forces (Jahan & Yumura, 2017; Nagasaki, Kanada et al., 2009). Out of the cells that assumed a bridge-shaped furrow, all wild type cells reached a cross-over point, defined as the point at which the bridge length equals the bridge width (Zhang & Robinson, 2005). In contrast, for *rnp1A* knockdown cells, only 6 out of 11 cells reached the cross-over point. The remaining 5 cells did not reach a cross-over point and assumed a morphology between the V-shape and bridge shape. The abnormality of *rnp1A* knockdown cell shape during cytokinesis implies lack of traction forces, which is typically caused by lack of adhesion (Jahan & Yumura, 2017).

Moreover, we measured the pole-to-pole distance after the cross-over point (**Fig. 1H**). For wild type cells, the pole-to-pole distance continued to increase after cells reached the cross-over point. However, for *rnp1A* knockdown cells, the pole-to-pole distance did not increase, and even decreased over time. The observed pole-to-pole distance difference reflects a defect in *rnp1A* knockdown cells ability to maintain cell-substrate adhesion (Poirier, Ng et al., 2012), and/or the ability to maintain normal microtubules (Snyder, Vogt et al., 1983; Giodini, Kallio et al., 2002). Notably, the RNP1A-OE cells did not show significant adhesion defects, and they did not show abnormal cell shape or pole-to-pole distance during cytokinesis. However, both *rnp1A* knockdown and RNP1A-OE cells showed slower growth in suspension (**Fig. 1H**), suggesting factors other than the identified defects in adhesion and cytokinesis also contribute to the slower cell growth.

### RNP1A enriches at the protrusive front, is slightly mechanoresponsive and interacts with IQGAP1 *in vivo*

To better understand how RNP1A exerts its functions in cells and how it interacts with CK proteins, we then investigated the intracellular localization of RNP1A. Using GFP-tagged RNP1A, we investigated the localization of RNP1A during random migration in live cells (**Fig. 2A**; **Movies EV 4A, 4B**). RNP1A-GFP did not localize to the protrusive front when the protrusion was initially formed. However, RNP1A-GFP enriched at the protrusive front tens of seconds after protrusions were initiated. We observed that sometimes the protrusion retracted after RNP1A-GFP enrichment. At other times, new protrusions formed on top of the RNP1A-GFP enriched protrusion. This observation resembles what was previously described where RNP1A localizes to protrusive front during chemotaxis (Ngo et al., 2016). The GFP-only control did not exhibit any enrichment to the protrusive front (**Fig. 2A; Movie EV 3**). To validate our live-cell observations, we performed immunofluorescence in fixed wild type parental cells (orfJ; Ax3(Rep orf+)) using anti-RNP1A antibodies. We observed that RNP1A enriched at protrusive region of the cell (**Fig. 2B**), which resembles what we observed with RNP1A-GFP in live cells.

**Figure 2.**
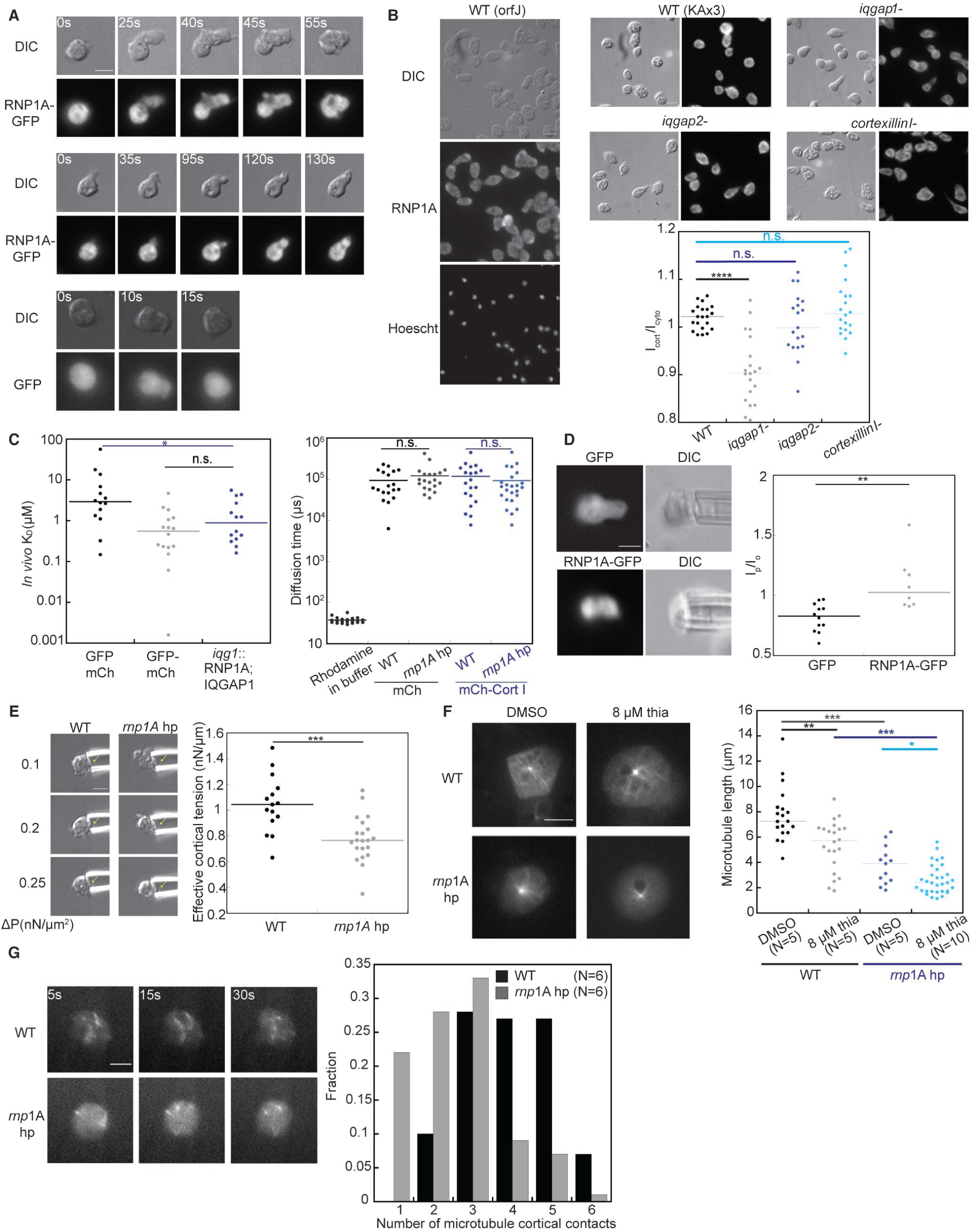
RNP1A localizes to the protrusive front of migrating cells, is slightly mechanoresponsive, interacts with IQGAP1 *in vivo*, and is important for microtubule polymer stabilization. (A) Images show localization of GFP and RNP1A-GFP during random cell migration. Scale bar, 10 µm. (B) Left: Immunofluorescence against RNP1A on fixed wild type cells (orfJ). Cells were fixed with -20°C acetone. Right: Immunofluorescence against RNP1A in wild type (KAx3), *iqgap1* null, *iqgap2* null and *cortexillin I* null cells. Cells were fixed with -20°C acetone. Quantification of cortical enrichment of RNP1A: cortex was manually traced for each single cell and mean intensity was measured for both cortical region and cytosolic region. Then the ratio of cortical to cytoplasmic intensity was taken as I_cort_/ I_cyto_. Data were collected from 20 wild type (KAx3), 19 *iqgap1* null, 18 *iqgap2* null, and 19 *cortexillin I* null cells. Scale bar, 10 µm. (C) Left: *In vivo* K_D_ between unlinked GFP and mCherry (negative control), GFP linked mCherry (positive control), as well as RNP1A-GFP and mCh-IQGAP1 were measured by FCCS. Data were collected from 13 cells expressing unlinked GFP and mCherry, 15 cells expressing GFP linked mCherry and 13 cells expressing RNP1A-GFP and mCh-IQGAP1. *In vivo* K_D_ between RNP1A-GFP and mCh-IQGAP1 was measured to be 0.89 µM (median). Right: Diffusion time of mCherry and mCh-Cortexillin I was quantified by FCS in either wild type control or *rnp1A* knockdown cells. Data were pooled from 18 wild type control cells expressing mCherry, 18 *rnp1A* knockdown cells expressing mCherry, 19 wild type control cells expressing mCh-Cortexillin I, 22 *rnp1A* knockdown cells expressing mCh-Cortexillin I, and 20 individual measurements of Rhodamine in imaging buffer. (D) Right: images show micropipette aspiration of cells expressing GFP or RNP1A-GFP. Scale bar, 10 µm. The degree of mechanoresponsiveness of RNP1A-GFP was quantified as GFP intensity ratio I_p_/I_o_, the ratio of mean signal intensity inside to outside of the pipette. Data were pooled from 11 wild type cells (orfJ) expressing GFP and 11 wild type cells (orfJ) expressing RNP1A-GFP. (E) Left: images show micropipette aspiration of wild type control and *rnp1A* knockdown cells at select applied negative pressure. Scale bar, 10 µm. Yellow arrows indicate portion of the cell inside the micropipette. Effective cortical tension was quantified from micropipette aspiration movies. Data were quantified from 14 wild type control cells and 20 *rnp1A* knockdown cells. (F) Wild type control or *rnp1A* knockdown cells expressing GFP-tubulin were treated with either 8 µM thiabendazole or equal volume of DMSO vehicle, and put under a thin 2% agarose sheet for better visualization. Scale bar, 10 µm. Microtubule lengths were quantified manually. Data were pooled from 5 wild type control cells treated with DMSO, 5 wild type control cells treated with 8 µM Thiabendazole, 5 *rnp1A* knockdown cells treated with DMSO, and 10 *rnp1A* knockdown cells treated with 8 µM Thiabendazole. Each dot represents the measurement from a single microtubule filament. (G) TIRF images of the bottom of wild type control and *rnp1A* knockdown cells expressing GFP-tubulin are shown. Scale bar, 5 µm. Number of microtubule contacts were manually counted for the duration of a 30-seconds movie for each cell at each second and summarized in the histogram. Data were pooled from measurements from 6 wild type control cells and 6 *rnp1A* knockdown cells. Data information: All statistical analysis was done with Kruskal–Wallis followed by Wilcoxon-Mann–Whitney test. *, P≤0.05; **, P≤0.01; ***, P≤0.001; ****, P≤0.0001; n.s, not significant.

Next, we investigated RNP1A localization under different CK protein null backgrounds to test if any CK protein is required for RNP1A localization. Interestingly, we observed that RNP1A is more cortically enriched in all KAx3 wild type and null mutants than in the orfJ background, highlighting a difference in the parental strains (**Fig. 2B)**. Out of all mutants, the *iqgap1* null mutant exhibited less RNP1A cortical enrichment than in the wild type and other strains, indicating that IQGAP1 helps facilitate RNP1A cortical localization (**Fig. 2B)**. This observation prompted us to further elucidate the interaction between RNP1A and IQGAP1. To do so, we employed fluorescence cross correlation spectroscopy (FCCS) to determine the *in vivo* K_D_ between RNP1A and IQGAP1 in the cytosol, and found that RNP1A and IQGAP1 interact *in vivo* with an apparent K_D_ of 0.88 μM (**Fig. 2C, left**). These observations indicate that IQGAP1 helps RNP1A localization by interacting with RNP1A. However, whether this interaction is direct or facilitated through other CK proteins such as Cortexillin I could not be concluded from FCCS data. Since IQGAP2, another key signaling protein in the CKs, also interacts with Cortexillin I (Kothari et al., 2019), we attempted to perform FCCS between RNP1A and IQGAP2. However, it was extremely difficult to co-express fluorophore labeled RNP1A with IQGAP2, and therefore we could not acquire enough data to obtain conclusive results. RBPs have been proposed to act as protein-protein interaction hubs due to their multi-valency and presence of intrinsically disordered regions (Chen & Mayr, 2022). Therefore, we tested if knocking down *rnp1A* changes the overall size of Cortexillin I-containing CKs by comparing the *in vivo* diffusion time of Cortexillin I in wild type control and *rnp1A* knockdown cell lines as a proxy. We observed no significant difference between wild type control cell line and *rnp1A* knockdown cell line in regard to diffusion time of mCherry-labeled Cortexillin I, suggesting that reduction of RNP1A might not significantly change the overall size of Cortexillin I-containing complexes (**Fig. 2C, right**). It remains possible that completely knocking out *rnp1A* might impact the size of the CKs, but this was not testable given the essential nature of *rnp1A*.

Proteins within the mechanoresponsive CKs, including IQGAP2, localize to the cleavage furrow during cytokinesis and to the posterior end of migrating cells, where Myosin II actively assembles to generate contractile forces (Srivastava & Robinson, 2015; Revington, McCloskey et al., 1987). On the other hand, IQGAP1, a component of the non-mechanoresponsive CKs, localizes to the front end of migrating cells (Brandt, Marion et al., 2007). Given that IQGAP1 interacts with RNP1A and regulates its localization, and the prior observation that RNP1A also enriches at the protrusive front during cell migration, we next assessed if RNP1A is, similarly to IQGAP1, non-mechanoresponsive. We performed micropipette aspiration (MPA) on wild type cells (orfJ) expressing GFP or RNP1A-GFP. Interestingly, we observed RNP1A-GFP was slightly mechanoresponsive, indicated by the increased ratio of GFP intensity in the micropipette compared to outside of the micropipette (**Fig. 2D**). This might not be surprising given that computational analysis has indicated the presence of what we term as ambiguous CKs, which contain both IQGAP1 and IQGAP2 (Plaza et al.; manuscript under review). Therefore, the possibility that RNP1A interacts with other mechanoresponsive CK proteins cannot be ruled out.

The observation of RNP1A interacting with CK proteins and its mechanoresponsive prompted us to test if RNP1A impacts cellular mechanics. To do so, we used MPA to measure effective cortical tension in wild type control and *rnp1A* knockdown cells, and observed a more than 20% reduction in effective cortical tension in *rnp1A* knockdown cells (**Fig. 2E**). Coincidentally, we also observed that *rnp1A* knockdown reduced cortical F-actin levels (**Fig. EV1**). Since the cortical actomyosin network architecture is important for optimal cortical tension (Chugh, Clark et al., 2017; Chugh & Paluch, 2018; Srivastava & Robinson, 2015; Luo, Srivastava et al., 2014), the reduced cortical F-actin amount may be a contributor to the decreased cortical tension in *rnp1A* knockdown cells.

### RNP1A is important for microtubule polymer stabilization and microtubule cortical contacts

Apart from the connection between RNP1A and the CKs proteins, we also previously discovered RNP1A functions in another set of important cytoskeletal structures that regulates cell shape, the microtubules. We identified RNP1A as a genetic suppressor of nocodazole, and over-expression of RNP1A protects microtubules from nocodazole (Ngo et al., 2016). Using the *rnp1A* hairpin knockdown cells, we tested if *rnp1A* knockdown sensitized microtubules to thiabendazole, a microtubule polymerization inhibitor. We measured microtubule length in wild type control cells and *rnp1A* knockdown cells under 0 μM (equal volume of DMSO vehicle) and 8 μM thiabendazole, which were put under a thin agarose sheet for better visualization (**Fig. 2E**). Treatment with 8 μM thiabendazole significantly decreased microtubule lengths in wild type cells. Under both 0 μM and 8 μM thiabendazole treatment, *rnp1A* knockdown cells had reduced microtubule lengths as compared to wild type cells (**Fig. 2E**). Therefore, *rnp1A* knockdown reduced microtubule lengths, further indicating that RNP1A is important for microtubule polymer assembly and/or stabilization.

Microtubule cortical contacts are important for signaling pathways that link to cortical actomyosin activity (Zhou et al., 2010). We then assessed how RNP1A contributes to microtubule dynamics. We measured the number of the microtubule cortical contacts in wild type and *rnp1A* knockdown cells through 30-second TIRF movies (**Fig. 2F; Movies EV5, EV6**). As compared to wild type cells, the *rnp1A* knockdown cells had increased fractions of time points that exhibited 3 or fewer microtubule cortical contacts and reduced fractions of time points that exhibited 4-6 microtubule cortical contacts (**Fig. 2F**). This suggests that RNP1A is important for maintaining microtubule cortical contacts.

Interestingly, our previous study suggests that microtubule polymer stabilization is important for cortical mechanics (Zhou et al., 2010). Treatment with 10 μM nocodazole reduced cortical tension by 60% (Zhou et al., 2010). Thus, the destabilization of microtubule polymers in *rnp1A* knockdown cells might also contribute to their reduced cortical tension.

### RNP1A knockdown poises cells to shift away from the vegetative growth to a more developmental-like transcriptional profile

As we continued to explore the functions of RNP1A and its impact on cell behaviors, we wanted to understand how it impacts cellular state as a whole, which could be revealed by whole transcriptome level changes in *rnp1A* mutants. Additionally, since RNP1A is a predicted RNA-binding protein, it is plausible that it could be directly involved in regulation of gene expression (Kishore et al., 2010). Therefore, we performed RNA sequencing on *rnp1A* knockdown cell lines. We generated two pairs of wild type control cell lines and *rnp1A* knockdown cell lines from two independent transformations (we refer to these two transformations as R1 and R2) and performed RNA sequencing on both pairs of cell lines.

Overall, the *rnp1A* knockdown cell line from R1 exhibited more pronounced *rnp1A* knockdown (96%) compared to R2 (88%) (**Fig. 3D**). Interestingly, we also saw increased numbers of both significantly up- and down-regulated genes in *rnp1A* knockdown cells from R1 compared to that of R2 (**Fig. 3A and Fig. EV 2; Datasets EV1, EV2**), suggesting that RNP1A regulation of gene expression might be dependent on its concentration in cells. We then first looked at the down-regulated genes in *rnp1A* knockdown cells from R1 (**Fig. 3C**), since *rnp1A* knockdown is more significant in this strain. All molecular function, biological process and pathway analyses suggest that the most significantly reduced gene expression clustered around ribosomes, and the most significantly down-regulated biological pathway is translation, accordingly. Another significantly down-regulated biological pathway is cell adhesion, which aligns with the observation that *rnp1A* knockdown cells were less adhesive. More interestingly, we then compared significantly down-regulated genes from RNA-seq in the *rnp1A* knockdown cell lines with proteins that were identified as the macropinocytotic proteome (Journet, Klein et al., 2012). We found that 54% of significantly down-regulated genes (R1) encode proteins identified as part of the macropinocytotic proteome (**Appendix Table S1**). In contrast, only 16% of all protein-coding genes encode proteins within the macropinocytotic proteome (calculated from numbers supplied from Journet et al., 2012 and Dictybase). This significantly increased fraction (P<0.0001; comparison of proportions) suggests that *rnp1A* knockdown also led to reduced gene expression in the macropinocytotic pathways, which were not specifically identified by gene ontology or pathway analysis potentially due to incomplete annotation. Importantly, previous studies indicate that cells in the vegetative growth phase exhibit a greater expression of genes associated with translation and macropinocytosis compared to cells in the developmental stage (cup, stalk and spore cells) (Kin, Forbes et al., 2018). Therefore, the dual down-regulation of gene expression in translation and macropinocytosis from *rnp1A* knockdown cells might suggest these cells were poised to shift away from the vegetative growth phase and transition to a more developmental-like phase.

**Figure 3.**
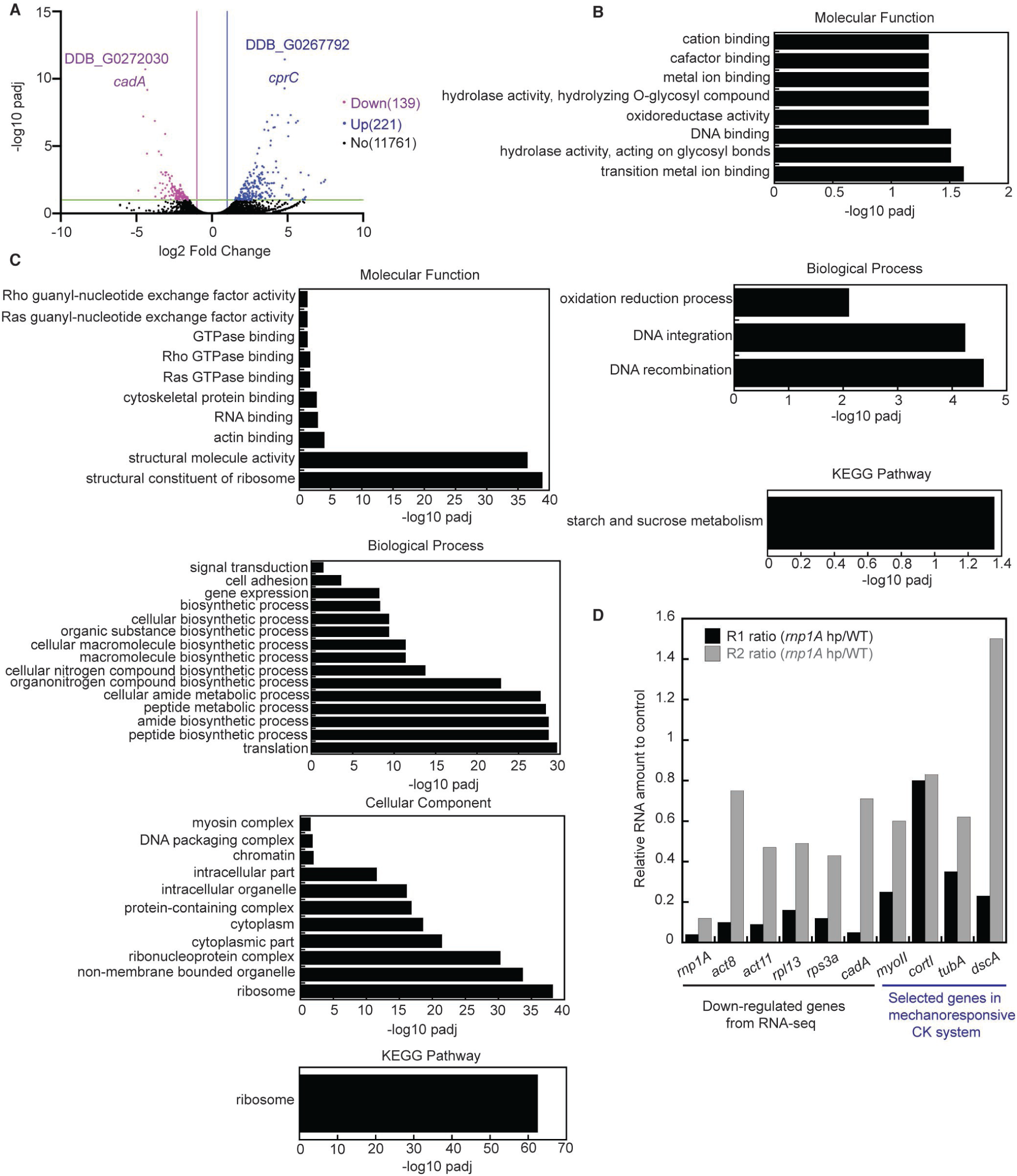
RNA-seq of *rnp1A* knockdown cells suggests that *rnp1A* knockdown reduces the expression of genes involved in translation and macropinocytosis. (A) Volcano plot of differentially expressed genes from RNA-seq of *rnp1A* knockdown cells is shown (replicate 1). Differentially expressed genes are at least two-fold up-regulated or down-regulated, and their corresponding padjs are equal or smaller than 0.1. Two most significantly up-regulated or down-regulated genes are labeled with DDB_ID or gene names. (B) Gene ontology analysis of up-regulated genes from RNA-seq of *rnp1A* knockdown cells (replicate 1). Gene ontology identification threshold is padj ≪ 0.05. (C) Gene ontology analysis of down-regulated genes from RNA-seq of *rnp1A* knockdown cells (replicate 1). Gene ontology identification threshold is padj ≪ 0.05. (D) Gene expression level of either selected significantly down-regulated genes or genes within the mechanoresponsive CK network compared to control cells from RNA-seq are shown for both replicate 1 and replicate 2.

We then looked at upregulated genes from RNA-seq data of *rnp1A* knockdown cells (R1). The most significantly enriched molecular function groups include hydrolase activity and oxidoreductase activity (**Fig. 3B**). Upon further investigation, genes clustered in the hydrolase activity group perform functions in carbohydrate catalysis (nagD, celA, alfA), which break down oligosaccharides (Dicytbase annotation; Blume & Ennis, 1991; Schopohl, Muller-Taubenberger et al., 1992). Genes clustered in the oxidoreductase activity group are majorly comprised of cytochrome P450 family genes (CYP556A1, CYP518B1, CYP513E1, CYP508A4, CYP508A3-2, CYP514A4). Although there are no specific annotations for functions of these genes in metabolism, cytochrome P450 family genes are generally considered to be involved in lipid and fatty acid metabolism (Chen, Capdevila et al., 2001; Van Bogaert, Groeneboer et al., 2011). Starch and sucrose metabolism was identified as a significantly up-regulated pathway (**Fig. 3B)**, which is one of the metabolic pathways that is more enriched in spore cells as compared to vegetative cells (Kin et al., 2018). Combining these metabolic features identified from up-regulated genes, we suspect that *rnp1A* knockdown cells to be actively metabolizing different biomolecules to maintain energy supply. Coincidentally, 53% of significantly upregulated genes in *rnp1A* knockdown cells overlap with genes enriched in cell types (cup, stalk and spore cells) in the development stage compared to cells in vegetative growth (Kin et al., 2018) (**Appendix Table S2; Dataset EV 3**). This 53% is notably higher than the 39% of all genes across the entire genome that become elevated during development (P<0.0001; comparison of proportion; calculated from numbers supplied from Kin et al., 2018 and Dictybase), again suggesting that these cells are poised to shift away from vegetative growth phase to a more development-like stage.

Next, we looked at selected genes involved in the mechanoresponsive CK system, including *myosin II, tubulin A, discoidin 1A*, and *cortexillin I*. Interestingly, almost all of these genes also showed down-regulation in *rnp1A* knockdown cell lines from both R1 and R2, except for *discoidin 1A* from R2 (**Fig. 3D**), although none of them qualify as significantly downregulated genes by statistical analysis in the context of the RNA-seq analysis (padj threshold ≪ 0.1). We also selected some significantly down-regulated genes (*act8, act11, rpl13, rps3a, cadA*) for comparison. In general, *rnp1A* knock down cells from R1 exhibited more pronounced down-regulation for all selected significantly down-regulated genes and selected genes involved in the mechanoresponsive CK system, suggesting a positive correlation between the degree of *rnp1A* knockdown and the degree of down-regulation of selected genes.

To validate the RNA-seq results, we first did qRT-PCR on select down-regulated genes from newly generated *rnp1A* knockdown cell lines (**Fig. 4A**). We observed fluctuations in *rnp1A* relative transcript level, and even more variations in relative transcript levels from some selected genes (*act8* and *act11, cadA, myoII, dscA*; *act8/11* primers recognize both *act8* and *act11* cDNA due to high degree of identity), suggesting variations from different biological replicates (**Fig. 4A**). However, it is not surprising given RNA-seq data from R1 and R2 revealed that a relatively small difference in *rnp1A* transcript level can lead to bigger differences in the transcript level of selected genes (**Fig. 3D**). Of genes that showed relatively smaller variations (*rpl13, rps3a, cortI, tubA*), median values suggest these genes showed reduced transcript level compared to wild type control (**Fig. 4A**), which agrees with the RNA-seq results.

**Figure 4.**
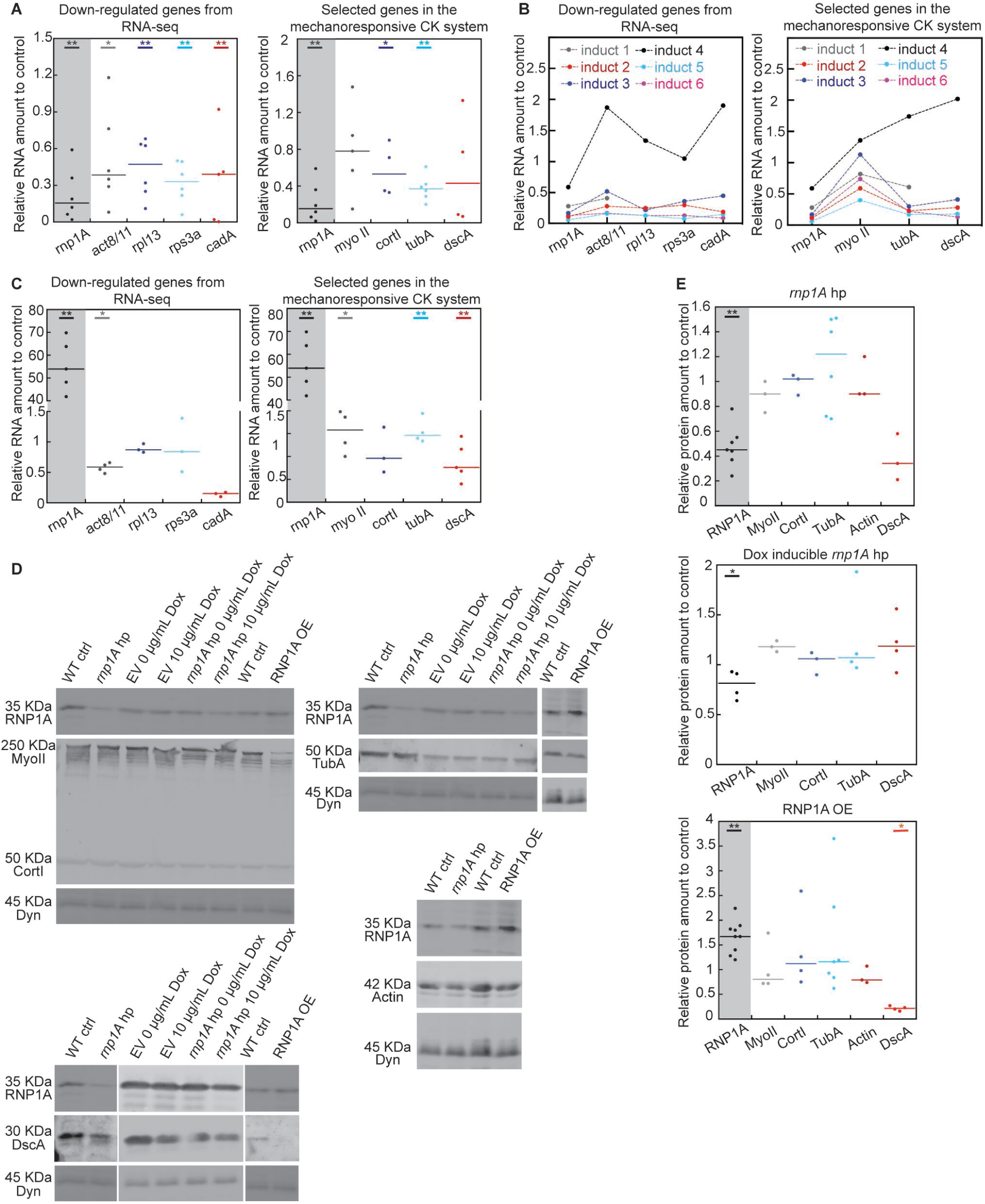
*rnp1A* knockdown, *rnp1A* doxycycline-inducible knockdown, and RNP1A overexpressing cells exhibit down-regulation of genes identified from RNA-seq by qRT-PCR; Discoidin 1A protein level is reduced in *rnp1A* overexpressing cells. (A) qRT-PCR validation on gene expression levels of selected genes identified from RNA-seq of *rnp1A* knockdown cells. Each dot represents the relative RNA amount determined from a single qRT-PCR run. Data were pooled from measurements of at least three biological replicates. (B) qRT-PCR validation on expression of selected genes in doxycycline-inducible *rnp1A* knockdown cells. Each dot represents a measurement from a single qRT-PCR run. Dots connected by dotted lines are from the same doxycycline induction. (C) qRT-PCR determined gene expression levels of selected genes in RNP1A overexpressing cells. Each dot represents the relative RNA amount determined from a single qRT-PCR run. Data were pooled from measurements from at least two biological replicates. (D) Western blots on Myosin II, Cortexillin I, Alpha Tubulin, Discoidin 1A in *rnp1A* knockdown, *rnp1A* doxycycline-inducible knockdown and RNP1A overexpressing cells. Dynacortin is a loading control. Western blot data were analyzed against total protein amount extracted from Coomassie staining of replica gels (**Appendix Fig.1**). (E) Quantification of protein amount of selected genes in CK network. Each dot represents the relative protein amount determined from a single blot. For measurement in *rnp1A* knockdown and overexpressing cells, data were pooled from measurements from at least two biological replicates. For measurement in doxycycline-inducible knockdown cells, data were pooled from at least three individual doxycycline induction. Data information: For (A), (C) and (E), all statistical analysis was done with Kruskal–Wallis followed by Wilcoxon-Mann–Whitney test. *, P≤0.05; **, P≤0.01; unlabeled, not significant.

All *rnp1A* knockdown cells used for RNA-seq were exposed to *rnp1A* hairpin for at least 3 weeks because we had to employ a slow-acting drug selection regimen to stabilize *rnp1A* knockdown. Therefore, these cells could have gone through extensive gene expression re-programming to adapt to the reduction of RNP1A. To monitor a more acute and first-line gene expression response to the reduction of RNP1A, we used the doxycycline-inducible system to induce the expression of the *rnp1A* RNA hairpin (**Fig. 4B**). In this system, *rnp1A* knockdown cells were exposed to *rnp1A* hairpin for only 48 or 100 hours. We observed doxycycline-inducible *rnp1A* knockdown also led to reduction in transcript levels for selected genes, except for one induction (induction 4) (**Fig. 4B**). We observed that the stronger the *rnp1A* knockdown level was, the more pronounced down regulation was for selected genes (**Fig. 4B**), again suggesting a positive correlation between degree of *rnp1A* knockdown and degree of down-regulation for these selected genes.

Additionally, we also examined gene expression changes in cells with overexpressed RNP1A (**Fig. 4C**). We found that *rpl13* and *rps3a* gene expression remained unchanged (**Fig. 4C**). Surprisingly, we found that *act8*/*act11* and *dscA* transcript levels were reduced as compared to wild type control (**Fig. 4C**), showing similar trends as in the *rnp1A* knockdown cells. On the other hand, the *myoII* and *tubA* transcript levels showed slight increases.

Finally, we examined protein level changes for select key CK system genes (**Figs. 4D, 4E; Appendix Fig. 1**). First, we observed that in doxycycline-inducible system, RNP1A protein knockdown level (20%; median) was not as pronounced as in persistent knockdown (52%; median) (**Figs. 4D, 4E**). Second, the only significant protein level reduction was observed for Discoidin 1A in RNP1A-OE cells. At the transcript level, *cortI* and *tubA* showed down-regulation from RNA-seq and qRT-PCR data in *rnp1A* knockdown cells, but their corresponding protein products were not significantly altered in expression (**Figs. 4D, 4E**). This reveals that transcript level expression change and protein level expression change do not always correlate.

### RNP1A binds to RNA transcripts encoding for proteins identified in the macropinocytotic proteome

Given that RNP1A is predicted to be an RBP, we next assessed which transcripts RNP1A binds to and how these might participate in the functional roles of RNP1A. We first identified the mRNA transcripts bound by RNP1A using CLIP-seq. Previously, it was discovered that RNP1A binds to ncRNA that regulates *Dictyostelium* development (Avesson, Schumacher et al., 2011), validating RNP1A as an RBP and revealing that RNP1A binds to non-messenger RNAs. Here, we were primarily interested in transcripts that have protein coding potential.

From the CLIP-seq analysis, we identified 24 mRNA transcripts that bind to RNP1A with a stringency beyond 90% confidence (padj ≪ 0.1) (**Fig. 5A**). Out of the 24 transcripts identified, one (DDB_G0283281, strictosidine synthase family protein) was also identified as significantly up-regulated by RNA-seq in both *rnp1A* knockdown cell lines from R1 and R2. More interestingly, we observed that some molecular functions of the genes coding for these 24 transcripts overlap with part of molecular functions of genes that were upregulated in *rnp1A* knockdown cells, including hydrolase activities and metal ion binding (**Figs. 3B, 5B**). Although genes in these overlapping functional groups were denoted with the same annotations, they are not the same set of genes, suggesting RNP1A might not directly regulate the abundance of the transcripts it binds to.

**Figure 5.**
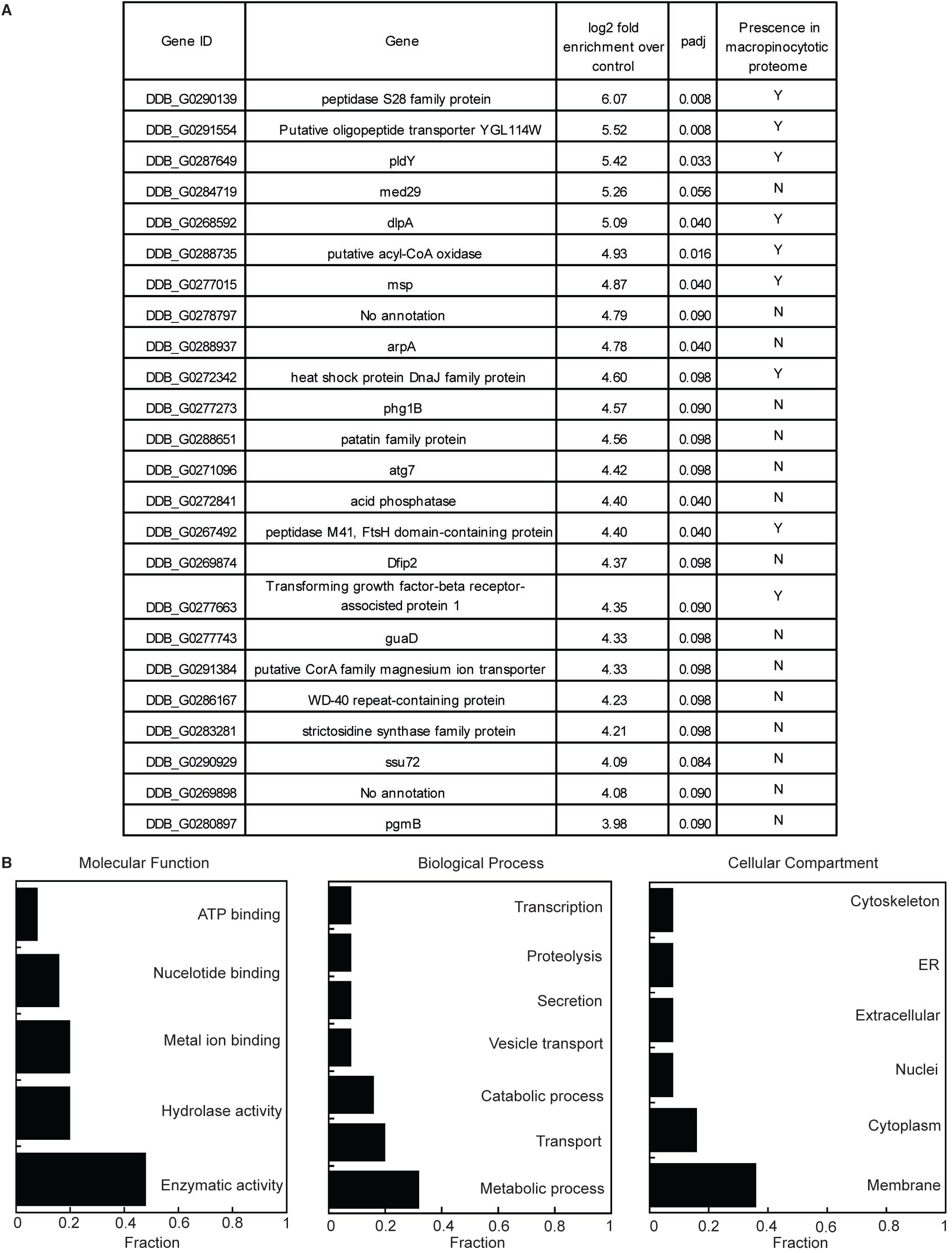
CLIP-seq revealed mRNA transcripts bound to RNP1A. (A) Full list of identity of mRNA transcripts bound to RNP1A-GFP, and their presence in the macropinocytotic proteome. “Y” = present in macropinocytotic proteome. “N” = not present in macropinocytotic proteome. Another transcript, WD repeat-containing protein 70 (DDB_G0283495; padj = 0.102) is not included in the list shown in figure due to its padj being slightly higher than 0.1. (B) Annotation of mRNA transcripts bound by RNP1A-GFP grouped by molecular functions, biological processes, and cellular components, and presented as fraction of all 24 transcripts identified to be bound to RNP1A.

Importantly, we noticed that 9 out of the 24 transcripts (37.5%) encode proteins within the macropinocytotic proteome (Journet et al., 2012) (**Fig. 5A**), which is significantly more enriched than the 16% of all protein-coding genes that encode proteins within the macropinocytotic proteome (P=0.0025; comparison of proportions; numbers supplied from Journet et al., 2012 and Dictybase). Out of these 9 transcripts encoded proteins, DlpA has been found to be important for cytokinesis, and *dlpA* null mutants have cytokinetic defects (Masud Rana et al., 2013). Moreover, DlpA localizes to the phagocytic cup during phagocytosis (Fujimoto, Tanaka et al., 2019), an endocytic process sharing same evolutionary origin and similar biochemical pathways with macropinocytosis (Vines & King, 2019). Furthermore, peptidase S28 family protein, putative acyl-CoA oxidase, pldY, peptidase M41, FtsH domain-containing protein, and msp are annotated with metabolic functions on different biomolecules. Although there is no direct evidence describing their roles in macropinocytosis apart from their association with the macropinocytotic proteome, it is likely they could be involved in digesting nutrients during macropinosome maturation (Vines & King, 2019). Moreover, many mechanoresponsive network proteins (RNP1A, Myosin II, IQGAP1, IQGAP2, Cortexillin I, Tubulin beta chain, Discoidin 1A) have also been identified within the macropinocytotic proteome, prompting the possibility that RNP1A might work together with some of these CK proteins during macropinocytosis.

### RNP1A contributes to macropinocytosis through regulation of transcripts it binds to and in coordination with Contractility Kit proteins

Macropinocytosis is the major pathway for *Dictyostelium* cells to uptake nutrients in fluid phase growth (Hacker, Albrecht et al., 1997; Williams & Kay, 2018). When macropinocytosis is reduced, the cells starve, which leads to initiation of a developmental program (Souza, da Silva et al., 1999) and suppression of gene expression in macropinocytosis and translation (Kin et al., 2018), as we observed from RNA-seq. One possible explanation for our RNA-seq and CLIP-seq data discussed above could be that RNP1A regulation on transcripts it binds to was diminished in *rnp1A* knockdown cells, which in turn leads to reduction of macropinocytosis and therefore starvation. Alternatively, RNP1A could have a direct impact on gene expression in translation and macropinocytosis regardless of its binding to transcripts. In either scenario, we hypothesized a decrease in macropinocytosis.

To test this, we measured cellular uptake of Tetramethylrhodamine isothiocyanate (TRITC)-labeled Dextran in wild type control and *rnp1A* knockdown cells (**Figs. 6A, 6B**). Indeed, *rnp1A* knockdown cells showed reduced uptake of TRITC-Dextran compared to wild type control. RNP1A-OE cells also showed reduced TRITC-Dextran uptake, although the reduction was less severe than in the *rnp1A* knockdown cells (**Fig. 6A**). The reduction in macropinocytosis from both *rnp1A* knockdown and overexpressing cells could explain slower growth rates in both mutants as discussed above, as cell proliferation is limited by nutrient uptake. We next asked if the defect in uptake of TRITC-Dextran is due to aberrant macropinocytotic crown formation. We measured the macropinocytotic crown area in *rnp1A* knockdown and wild type control cells and found no significant difference (**Figs. 6C, 6D**). We then compared the number of macropinocytotic events per cell within a two-minute interval. As compared to wild type control, *rnp1A* knockdown cells had an increased fraction of cells that did not have any macropinocytotic events. Furthermore, no *rnp1A* knockdown cells had 3 or 4 macropinocytotic events, whereas wild type control cells did (**Fig. 6D**). Overall, the defect in TRITC-Dextran uptake in *rnp1A* knockdown cells is due to decreased macropinocytosis frequency. We next examined RNP1A localization during macropinocytosis (**Fig. 9A**). GFP-tagged RNP1A signal was slightly enriched around the macropinosome formation site between macropinosome formation and early macropinosome retraction into the cell body, whereas GFP control did not show any enrichment (**Fig. 9A; Movies EV7, EV8**).

**Figure 6.**
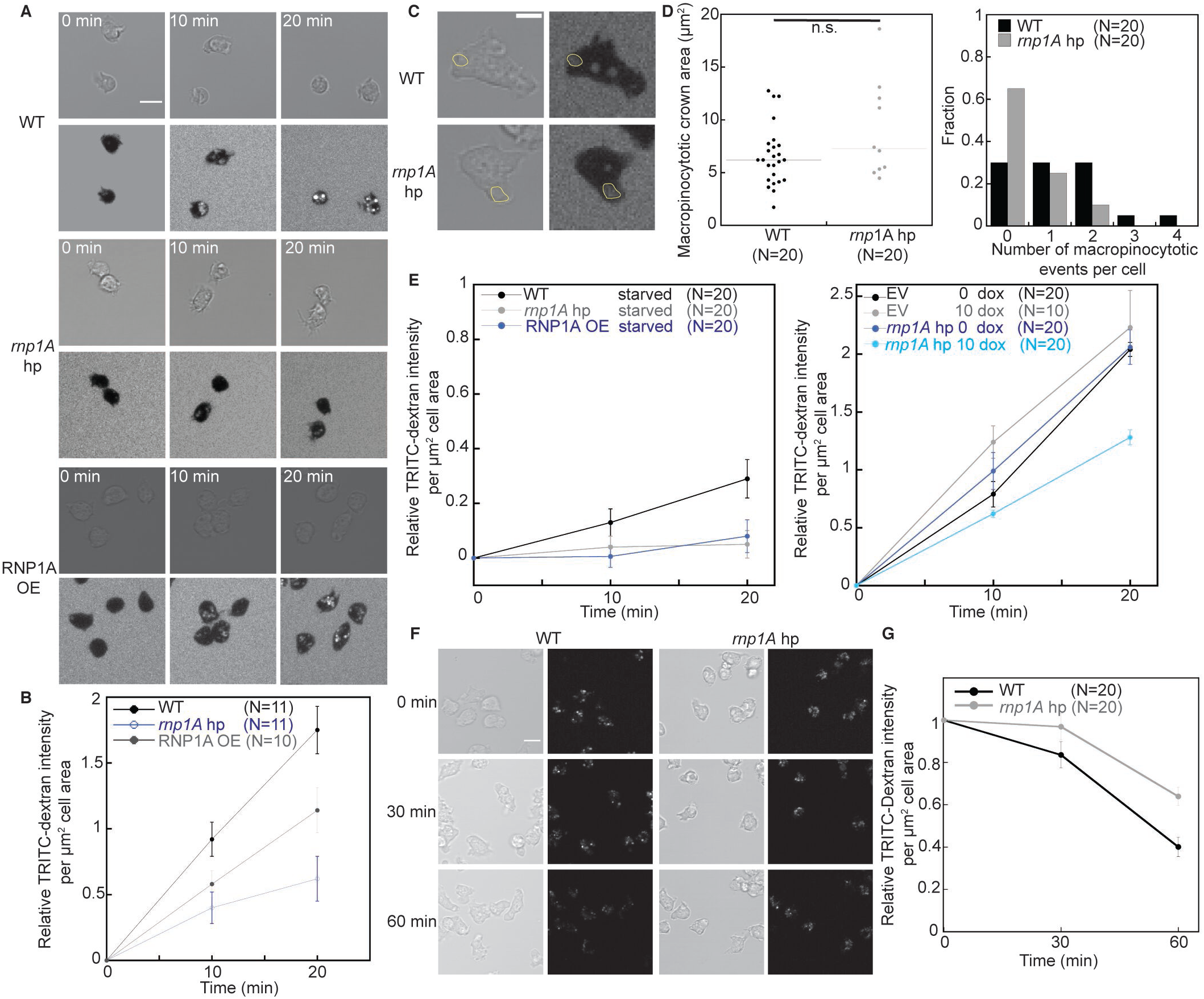
*rnp1A* knockdown cells show defects in macropinocytosis. (A) and (B) TRITC-Dextran uptake in *rnp1A* knockdown and RNP1A overexpressing cells over the span of 20 minutes. TRITC-Dextran intensity was quantified by mean TRITC intensity of each cell, background subtracted, normalized to cell area, and then normalized to that at the first time point. Scale bar, 10 µm. Data were pooled from 11 wild type control cells, 11 *rnp1A* knockdown cells, and 10 RNP1A overexpressing cells. Error bars indicate standard errors. (C) and (D) Quantification of macropinocytotic crown area and number of macropinocytotic events per cell over the span of 2 minutes. Macropinocytotic crowns were manually traced as shown in yellow circles in images at the first time point of crown membrane closure. Scale bar, 5 µm. Data were pooled from 20 wild type control cells and 20 *rnp1A* knockdown cells. (E) TRITC-Dextran uptake in starved *rnp1A* knockdown, RNP1A overexpressing cells, and *rnp1A* doxycycline-inducible knockdown cells. Data were pooled from 10 or 20 cells as indicated in the figure. Error bars indicate standard errors. (F) and (G) Panel shows loss of internalized TRITC-Dextran signal in wild type control and *rnp1A* knockdown cells over the span of 60 minutes. Scale bar, 10 µm. TRITC-Dextran intensity was quantified by mean TRITC intensity of each cell, background subtracted, normalized to cell area, and then normalized to that at first time point. Error bars indicate standard errors. Data were pooled from 20 wild type control cells and 20 *rnp1A* knockdown cells. Data information: For (D), statistical analysis was done with Kruskal–Wallis followed by Wilcoxon-Mann– Whitney test. n.s., not significant.

Starvation slows down macropinocytosis, including internalization and transfer of material between endocytic compartments (Smith, Lima et al., 2010), and this could be a reason for the reduction in macropinocytosis frequency observed in *rnp1A* knockdown cells. We confirmed that in all wild type control, *rnp1A* knockdown and overexpressing cells, TRITC-Dextran uptake is dramatically slowed down after two hours of starvation (**Fig. 6E; Appendix Fig. 2**). Therefore, we suspect that long-term *rnp1A* knockdown may lead to long-term starvation, and therefore exacerbates the reduction in TRITC-Dextran uptake we observed (**Figs. 6A, 6B**). To test this, we used doxycycline-inducible *rnp1A* knockdown cells to test if acute *rnp1A* knockdown led to similar or lesser degree of reduction in macropinocytosis. Indeed, we still saw reduction in macropinocytosis in the doxycycline-inducible *rnp1A* knockdown cells, though not as severe as in persistent *rnp1A* knockdown cells (**Fig. 6E; Appendix Fig. 2**).

We next asked if the reduction in macropinocytosis is partially regulated by transcripts to which RNP1A binds. We first validated if proteins encoded by RNP1A-binding transcripts are indeed involved in macropinocytosis. To do so, we picked *dlpA* to study because DlpA showed the highest spectral count (116) in the mass spectrometry of macropinocytotic proteome (Journet et al., 2012) out of the 9 proteins encoded by RNP1A-binding RNAs. To visualize localization of DlpA during macropinocytosis, we expressed GFP-tagged DlpA in wild type cells. Strikingly, DlpA enriched at the macropinocytotic crown and continued to encircle the macropinosome after formation and internalization (**Fig. 9B; Movie EV 9**). Then, we asked if loss of the *dlpA* could also lead to macropinocytosis defect. To do so, we used *dlpA* null cells (Miyagishima, Kuwayama et al., 2008). Interestingly, we did not see a reduction in TRITC-Dextran uptake in *dlpA* null cells compared to wild type parental cells (Ax2) (**Fig. 7A**). However, when we measured the rate of loss of internalized TRITC-Dextran, we did observe a reduction in the rate of loss of internalized TRITC-Dextran in *dlpA* null cells compared to wild type parental cells (Ax2) (**Fig. 7B**). This result indicates that DlpA might be involved in macropinosome maturation when nutrients in macropinosomes are degraded for cellular use. Newly formed macropinosomes do not mature until after around 60 seconds, when V-ATPase is delivered to macropinosomes (Cardelli, 2001). Therefore, we suspect that DlpA involvement with macropinosomes might lapse over 60 seconds. Indeed, we observed that GFP-DlpA signal encircling internalized TRITC-Dextran persist for around 159 s (median; SE=12.9 s; N=8) (**Fig. 7C**), again suggesting that DlpA function in macropinocytosis might be involved in macropinosome maturation.

**Figure 7.**
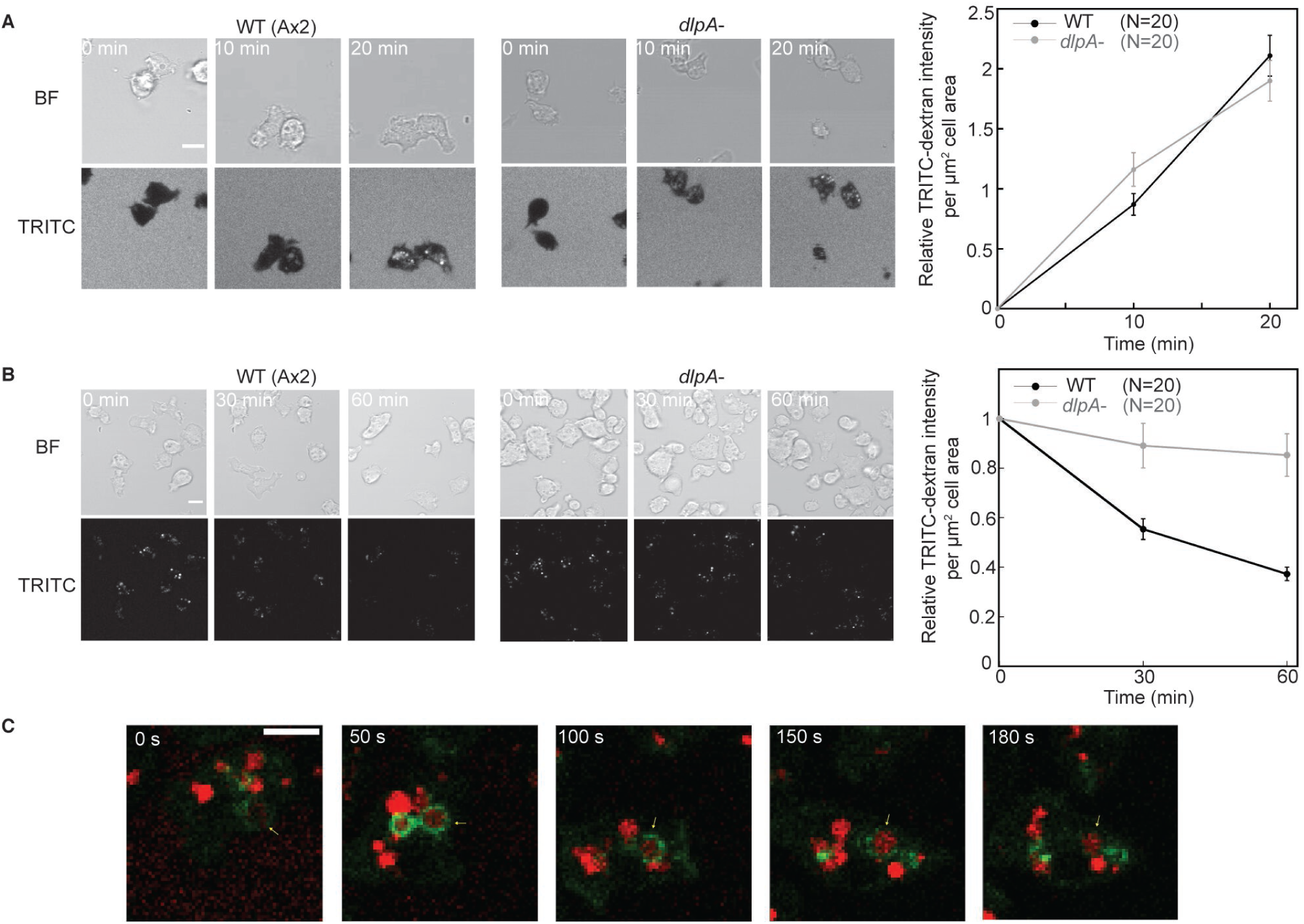
*dlpA* null cells show delayed loss of internalized TRITC-Dextran, and GFP-DlpA persists around macropinosomes for over 150 seconds. (A) TRITC-Dextran uptake in wild type (Ax2) and *dlpA* null cells over the span of 20 minutes. TRITC-Dextran intensity was quantified by mean TRITC intensity of each cell, background subtracted, normalized to cell area, and then normalized to that at the first time point. Scale bar, 5 µm. Data were pooled from 20 wild type cells (Ax2) and 20 *dlpA* null cells. Error bars indicate standard errors. (B) Internalized TRITC-Dextran degradation in wild type (Ax2) and *dlpA*-cells over the span of 60 minutes. TRITC-Dextran intensity was quantified by mean TRITC intensity of each cell, background subtracted, normalized to cell area, and then normalized to that at first time point. Scale bar, 10 µm. Data were pooled from 20 wild type cells (Ax2) and 20 *dlpA* null cells. Error bars indicate standard errors. (C) GFP-DlpA localization around macropinosome during after internalization of TRITC-Dextran. Green: GFP-DlpA; red: TRICT-Dextran. Scale bar, 10 µm. Yellow arrows point to the same macropinosome over time.

To test if *dlpA* transcript could be regulated by RNP1A at all, we investigated if *rnp1A* knockdown cells might partially phenocopy *dlpA* null cells during macropinocytosis. Therefore, we assessed TRITC-Dextran loss in *rnp1A* knockdown cells, and indeed, we observed a reduction of rate in the loss of internalized TRITC-Dextran in *rnp1A* knockdown cells as compared to wild type control cells (**Figs. 6F, 6G**), similar to *dlpA* null cells. Thus, it is plausible that RNP1A might help regulate translation of *dlpA* transcript, and *rnp1A* knockdown cells partially present a macropinocytotic defect through lack of DlpA. This explanation does not exclude the possibility, however, that other genes down-regulated in *rnp1A* knockdown cells could also slow down macropinosome maturation, and therefore bypass RNP1A regulation of the transcripts to which RNP1A binds.

To test the hypothesis that RNP1A regulates transcripts to which it binds from an orthogonal approach, we investigated another transcript that is not involved in macropinocytosis, *atg7*. Previously, *atg7* was discovered to be involved in the autophagy pathway. Moreover, *atg7* null cells have fewer GFP-Atg18 (Otto, Wu et al., 2004) aggregates compared to wild type control cells (King, Veltman et al., 2011) when placed under mechanical stress. Therefore, we used doxycycline-inducible *rnp1A* knockdown cells expressing GFP-Atg18 to test if RNP1A regulates expression and/or function of *atg7*. When subjected under mechanical stress exerted by an agarose sheet, the doxycycline-induced *rnp1A* knockdown cells had fewer GFP-Atg18 aggregates than the wild type control cells had (**Fig. EV3**), suggesting Atg7 requires RNP1A for its function and/or expression. Overall, these observations support the idea that RNP1A regulates the translational output of the transcripts and/or the function of the proteins encoded by the transcripts to which it binds.

Since we found RNP1A to interact with Cortexillin I and IQGAP1 *in vivo*, we next asked if these two proteins, along with other CK components, namely Myosin II and IQGAP2, contribute to macropinocytosis as well. We first assessed Cortexillin I, IQGAP1 and IQGAP2, since these mutant cells were all created from the same parental wild type strain (KAx3). Compared to wild type cells, *cortexillin I* null and *iqgap1* null cells both had decreased TRITC-Dextran uptake and smaller macropinocytotic crown area, whereas *iqgap2* null cells exhibited no defect in TRITC-Dextran uptake and crown size (**Figs. 8A, 8C**). None of these mutants showed significant differences in numbers of macropinocytotic events within a two-minute interval compared to wild type cells. Overall, the defect of macropinocytosis caused by loss of Cortexillin I and IQGAP1 was due to decreased macropinocytotic crown size, which is consistent with their roles in cell shape regulation, but distinct from what caused the macropinocytotic defect in *rnp1A* knockdown cells. We then assessed the localization of these proteins during macropinocytosis. We observed mCherry-Cortexillin I signal enriched at the cortical region around the macropinosome formation site (**Fig. 9C; Movie EV 10**). The GFP-IQGAP1 signal remained cortically localized and did not show distinct changes in localization, whereas the GFP-IQGAP2 signal only became slightly enriched at the tip of macropinocytotic crown after membrane closure (**Figs. 9D, 9E; Movies EV 11, 12**).

**Figure 8.**
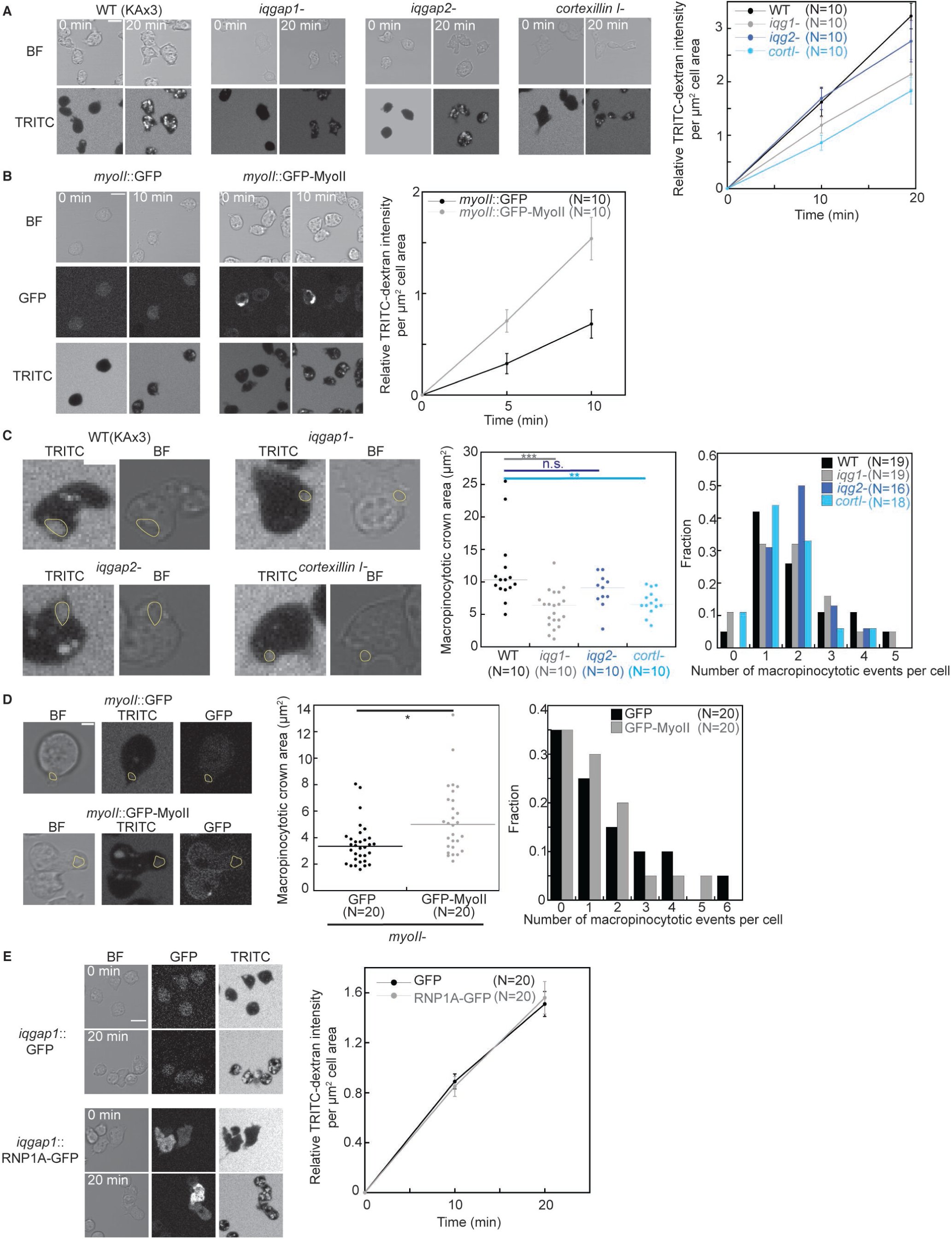
*iqgap1* null, *cortexillin I* null and *myosin II* null cells show defects in macropinocytosis. (A) TRITC-Dextran uptake in wild type (KAx3), *iqgap1*-, *iqgap2*- and *cortexillin I*-cells over the span of 20 minutes. TRITC-Dextran intensity was quantified by mean TRITC intensity of each cell, background subtracted, normalized to cell area, and then normalized to that at first time point. Scale bar, 10 µm. Error bars indicate standard errors. Data were pooled from 10 cells from each cell line. (B) TRITC-Dextran uptake in *myosin II* null cells expressing GFP or GFP-Myosin II over the span of 10 minutes. TRITC-Dextran intensity was quantified by mean TRITC intensity of each cell, background subtracted, normalized to cell area, and then normalized to that at first time point. Scale bar, 10 µm. Error bars indicate standard errors. Data were pooled from 10 cells from each cell line. (C) Quantification of macropinocytotic crown area and number of macropinocytotic events per cell over the span of 2 minutes. Macropinocytotic crowns were manually traced as shown in yellow circles in images. Scale bar, 5 µm. Data were pooled from the number of cells indicted in the figure. (D) Quantification of macropinocytotic crown area and number of macropinocytotic events per cell over the span of 2 minutes. Macropinocytotic crowns were manually traced as shown in yellow circles in images. Scale bar, 5 µm. Data were pooled from measurement from 20 cells per cell line. (E) TRITC-Dextran uptake in *iqgap1* null cells expressing GFP and RNP1A-GFP. TRITC-Dextran intensity was quantified by mean TRITC intensity of each cell, background subtracted, normalized to cell area, and then normalized to that at first time point. Scale bar, 10 µm. Error bars indicate standard errors. Data is pooled from 20 cells per cell line. Data information: For (C) and (D), all statistical analysis was performed with Kruskal–Wallis followed by Wilcoxon-Mann–Whitney test. *,P≤0.05; **,P≤0.01;***,P≤0.001; n.s, not significant.

**Figure 9.**
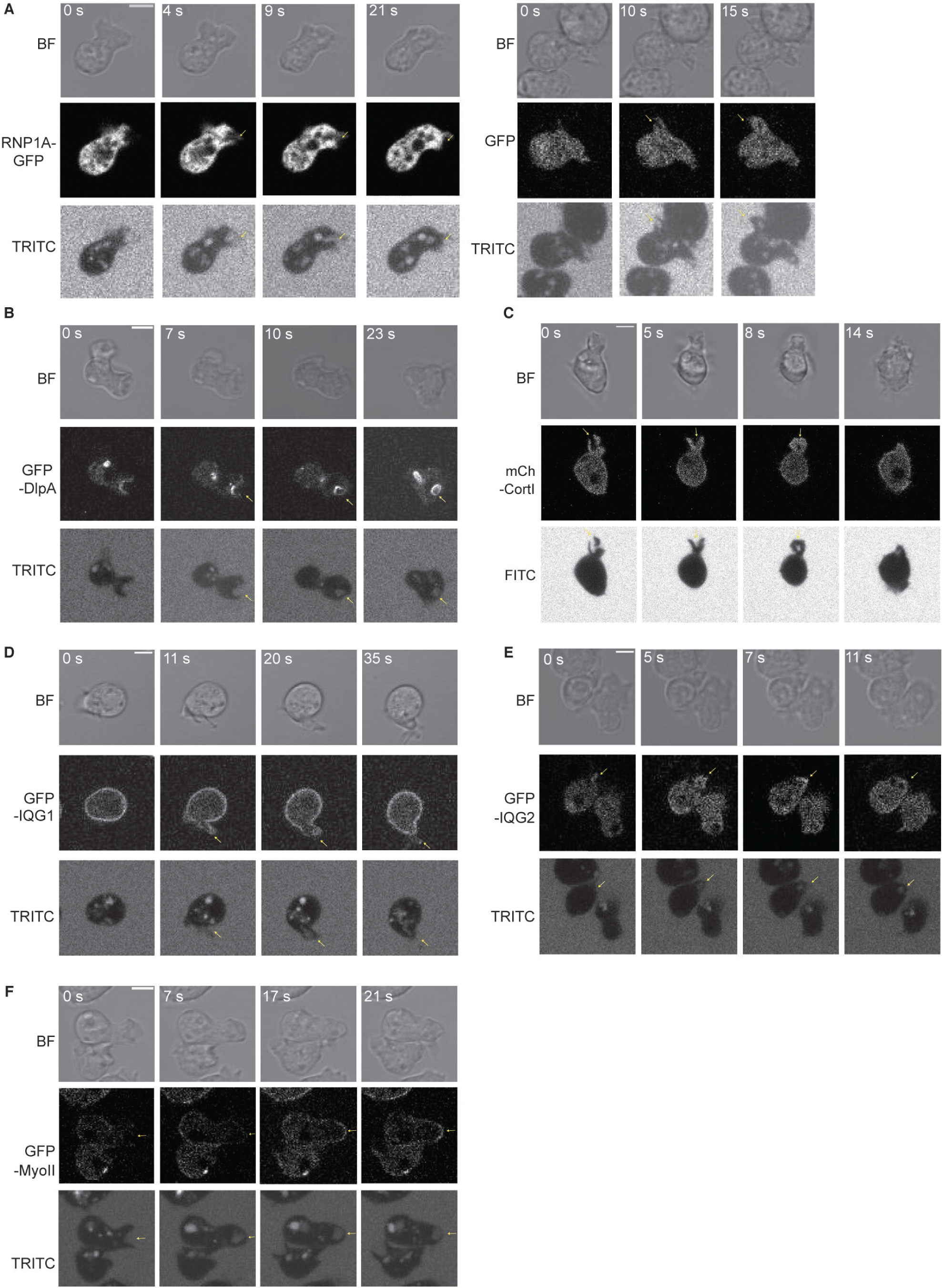
Localization of RNP1A, DlpA and CK proteins during macropinocytotic crown formation and closure. (A) Left panel shows localization of RNP1A-GFP during macropinocytotic crown formation and closure. Right panel shows localization of GFP during macropinocytotic crown formation and closure. (B) Panel shows GFP-DlpA localization during macropinocytotic crown formation and closure. (C) Panel shows mCherry-Cortexillin I localization during macropinocytotic crown formation and closure. (D) Panel shows GFP-IQGAP1 localization during macropinocytotic crown formation and closure. (E) Panel shows GFP-IQGAP2 localization during macropinocytotic crown formation and closure. (F) Panel shows GFP-Myosin II localization during macropinocytotic crown formation and closure. (A)-(F), yellow arrows point to sites of macropinocytotic crown formation. Scale bars, 10 µm.

We next explored if there is any crosstalk between RNP1A and IQGAP1 regulation of macropinocytosis, since IQGAP1 interacts with RNP1A *in vivo* and regulates RNP1A cortical localization. We observed no difference in RNP1A-GFP localization during macropinocytosis in *iqgap1* null cells compared to wild type control cells (**Appendix Fig. 3**). However, when we overexpressed RNP1A with exogenous RNP1A-GFP in *iqgap1* null cells, RNP1A overexpression did not lead to defects in macropinocytosis (**Fig. 8E**), as it did in wild type cells (**Figs. 6A, 6B**). This suggests that RNP1A overexpression works through IQGAP1 to cause macropinocytotic defects, suggesting a crosstalk between RNP1A and IQGAP1 during macropinocytosis.

Then, we assessed Myosin II’s involvement in macropinocytosis. We observed that compared to wild type cells (*myosin II*::GFP-Myosin II), *myosin II* null cells (*myosin II*::GFP) exhibited less TRITC-Dextran uptake and had a smaller macropinocytotic crown area (**Figs. 8B, 8D**). GFP-Myosin II signal also became enriched at the cortical region of macropinosome formation site after the macropinosome completely formed and started retracting back into the cell body (**Fig. 9F; Movie EV 13**). Moreover, both over-assembled (3xAla) or under-assembled (3xAsp) Myosin II (Hostetter, Rice et al., 2004) reduced TRITC-Dextran uptake, suggesting wild type Myosin II assembly dynamics help facilitate macropinocytosis (**Fig. EV4A**). Both mutants displayed reduction in macropinocytotic crown area compared to wild type cells (**Figs. 8D, EV 4B**). The 3xAsp-expressing mutant, however, exhibited a higher fraction of cells that had only one macropinocytotic event per cell, and none that had more than 3 macropinocytotic events per cell during a 2-minute interval (**Fig. EV 4B**). Thus, compared to wild type cells, the 3xAsp mutant was less efficient in performing macropinocytosis. Lastly, RNP1A-GFP overexpression did not cause macropinocytotic defects in *myosin II* null cells, similarly to in *iqgap1* null cells (**Figs. EV4A, 6A, 6B, 8E**). This suggests a crosstalk between RNP1A and Myosin II during macropinocytosis.

In the end, we observed that in *iqgap1* null, *cortexillin I* null, and *myosin II* null cells, GFP-DlpA localization during macropinocytosis remained the same as in wild type cells (**Appendix Fig. 4**), suggesting there might not be a direct a crosstalk between DlpA and CK proteins. Overall, we suspect that RNP1A works coordinately with CK proteins including IQGAP1 and Myosin II and through its binding to RNA transcripts such as *dlpA* to facilitate macropinocytosis.

## Discussion

Previous work has shown that RNA-binding proteins are involved in multiple biological processes (Gerstberger, Hafner et al., 2014; Hentze, Castello et al., 2018). In *Dictyostelium discoideum*, limited evidence suggests that RNA-binding proteins, such as Pumillo, are involved in localizing specific RNA transcripts to a sub-cellular location where the encoded protein functions (Hotz & Nelson, 2017). Other RBPs in *Dictyostelium discoideum* have been shown to function in post-transcriptional regulation (Bai, Wells et al., 2021), miRNA processing (Meier, Kruse et al., 2016), and miRNA maturation (Kruse, Meier et al., 2016). However, the function of RBPs within the mechanoresponsive system and relationship with the Contractility Kit proteins have not been explored before. Previously, we discovered that RNP1A interacts with Cortexillin I (Kothari et al., 2019) and is a genetic suppressor of nocodazole when over-expressed (Ngo et al., 2016). Here, we present that RNP1A is important for cell growth, adhesion, and normal cytokinesis. We discovered that RNP1A interacts with CK protein IQGAP1, is slightly mechanoresponsive, and contributes to cortical tension. RNP1A facilitates cortical microtubule contacts and stabilization of microtubule polymers. Furthermore, knockdown of *rnp1A* shifted cells away from vegetative growth to a more developmental stage-like transcriptional profile. RNP1A binds to transcripts that encode proteins involved in macropinocytosis and works alongside CK proteins to facilitate macropinocytosis. Based on all evidence gathered, we proposed a working model for the function of RNP1A (**Appendix Fig. 5**), suggesting its role in facilitating macropinocytosis, and ultimately promoting *Dictyostelium* vegetative growth.

We suspect the *rnp1A* knockdown phenotypes of growth and adhesion defects to be linked to gene expression changes. The alterations in the gene expression profile suggest a general shift away from a vegetative growth state in *rnp1A* knockdown cells, which is consistent with the reduced vegetative growth rate and macropinocytotic defects. Similarly, cell adhesion is found to be down-regulated through our RNA-seq data, and we observed adhesion defects in *rnp1A* knockdown cells as well. We also observed aberrant cytokinesis in *rnp1A* knockdown cells, but not in RNP1A-overexpressing cells. Morphologically, the inability to form the bridge-shaped cleavage furrow is similar to phenotype observed in cytokinesis A (Nagasaki et al., 2009), where cells depend majorly on constriction forces rather than traction forces to complete cytokinesis, reflecting a reduction in cell-substrate adhesion. We also observed that *rnp1A* knockdown and RNP1A overexpressing cells exhibit some similar trends phenotypically (growth, adhesion, gene expression and macropinocytosis), which may suggest that a wild-type level expression of RNP1A is required for optimal cell behaviors.

We have previously described *in vivo* interactions between RNP1A and Cortexillin I. Here, we discovered that RNP1A also interacts with IQGAP1, a component of the non-mechanoresponsive Contractility Kits. The localization of RNP1A does not exhibit the distinctive cortical localization of Cortexillin I (Cha & Jeon, 2011; Faix, Steinmetz et al., 1996) or IQGAP1 in orfJ (Ax3(Rep orf+) parental strain. RNP1A does have slightly enriched cortical distribution in KAx3 parental strain, suggesting an inherent difference exists between these parental strains. Interestingly, IQGAP1 has been shown to interact with active Rac1A (Faix, Clougherty et al., 1998), and Rac1A localizes to the macropinocytotic crowns (Williams, Paschke et al., 2019), which trends similarly with RNP1A localization during macropinocytosis. We were unable to achieve conclusive results on *in vivo* interaction between RNP1A and IQGAP2, a component of mechanoresponsive CKs. However, GFP-tagged RNP1A was slightly mechanoresponsive (slight cortical accumulation in response to imposed mechanical stress), consistent with RNP1A’s interactions with Cortexillin I, a mechanoresponsive CK protein.

RNA-binding proteins play a role in gene expression regulation via multiple mechanisms (Ray, Kazan et al., 2013). Polyadenylate-binding proteins (PABPs) have high sequence similarities with RNP1A (Ngo et al., 2016). PABPC helps regulate RNA polyadenylation, translation, localization and decay (Wigington, Williams et al., 2014). PABPC has also been identified to localize to the leading edge of migrating cells (Woods, Roberts et al., 2002), similar to RNP1A’s localization. Another class of proteins that have high sequence similarities with RNP1A are the cold-induced RNA-binding proteins, including Cirbp and Rbm3. These two proteins help regulate RNA translation and stability by controlling alternative polyadenylation (Liu, Hu et al., 2013). Rbm3 also localizes to invasive pseudopodia of mesenchymal breast cancer cells (Mardakheh, Paul et al., 2015) and regulates cell spreading and migration (Pilotte, Kiosses et al., 2018). Although characterized in different organisms, the similar localization patterns of PABPC and Rbm3 to RNP1A may suggest that there could be more functional similarities between this group of proteins.

What are the possible mechanisms by which RNP1A interacts with transcripts and proteins and how might these activities feed into RNA stability and/or protein translation? First, RNP1A is unlikely to regulate RNA turnover or decay during vegetative growth given that only one transcript to which RNP1A binds exhibited gene expression level changes. Instead, it is more likely that RNP1A regulates localization, translation, and/or modification of these transcripts during vegetative growth. Loss of RNP1A regulation on these transcripts could affect the translational output from these transcripts, leading to macropinocytotic defects, which we have observed for the *rnp1A* knockdown cells. Another, but not mutually exclusive, possibility is that RNP1A could have direct transcription factor ability, as it has been indicated that some RBPs can interact with chromatin to affect gene expression (Du & Xiao, 2020). While we cannot entirely rule out this possibility, we suspect this is unlikely since RNP1A did not show localization within the nucleus during vegetative growth. In regard to the possible role of RNP1A regulating translation, since localized translation takes place on the timescale of hundreds of seconds (Wang et al., 2018), it is less likely that RNP1A directs localized translation concurrently during macropinosome formation. Because macropinosome formation only takes about 5 -10 seconds from initiation to membrane closure and retraction completion. It is more likely that RNP1A regulation of its bound transcripts is an ongoing process in the cytoplasm. In this study, we did not directly test the posttranscriptional regulation by RNP1A on the transcripts to which it binds. Given the multi-layered functions of RNP1A, it is very challenging to disentangle mechanism crosstalks. We are, however, pointing out regulation of translational output as a possibility, which could warrant follow-up studies.

Macropinocytosis is a biological process where robust cell and cortical shape changes take place. Cell mechanics, including cortical tension effects, may help resist and then facilitate the progression of macropinocytosis. During the initiation stage of macropinocytosis, cortical tension is likely to resist macropinocytosis, and therefore active force production through actin polymer assembly and/or a relaxation of cortical tension may help promote macropinocytosis. In myoblasts for example, an acute decrease in plasma membrane tension results in initiation of macropinocytosis via phosphatidic acid production and PI(4,5)P_2_-enriched membrane ruffling (Loh, Chuang et al., 2019). In the *Hydra vulgaris* outer epithelial layer, mechanical stretch inhibits macropinocytosis via activating stretch-activated channels, including Piezo, leading to calcium influx (Skokan, Hobmayer et al., 2021). After initiation of macropinocytosis, the macropinocytotic crown membrane continues to elongate and eventually closes up. Once elongated, cortical tension along with active inward force production can drive the pull back of the cortex, re-rounding the cell. Myosin proteins play several roles in this process. For example, some Myosin I isoforms are recruited to the macropinocytotic crowns/cups and form a broad ring around the cortex (Brzeska, Koech et al., 2016). Myosin II has been implicated in Neuro-2a cells where Myosin IIB is essential for macropinocytosis (Jiang, Kolpak et al., 2010). We found that in *Dictyostelium discoideum*, Myosin II and its assembly dynamics are essential for normal macropinocytosis. Myosin II localizes to the cortex around the macropinocytotic crown where it helps drive macropinosome retraction after crown closure. Moreover, the involvement of CK proteins in macropinocytosis indicate that cortical mechanics and mechanoresponse feed into macropinocytosis. The transient localization of IQGAP2 right after crown closure likely signifies the transition from macropinocytotic crown formation to retraction, helping to recruit Myosin II to the crown to help drive retraction.

Our studies also raise an interesting question: Do mechanical stimuli affect RNA regulation in cells, including, but not limited to, RNA localization, translation, processing, decay, and/or modification, and vice versa? Although this study does not directly address this connection, the association between RNP1A with the CK machinery, the mRNAs it binds, and the association between all these factors with cell shape changes strongly support some level of interconnection between these processes. Some budding evidence from the literature also supports such a connection, suggesting this interconnection will likely turn out to be a much more fundamental concept than is currently appreciated. For example, mechanical stimuli can recruit polyA mRNA and ribosomes via focal adhesion and associated proteins (Chicurel, Singer et al., 1998). More recently, heterogeneous nuclear ribonucleoprotein C (hnRNPC) was observed to localize to the cardiomyocyte sarcomeres, and ECM remodeling in pathological conditions leads to hnRNPC association with the translational machinery. Furthermore, hnRNPC then regulates the alternative splicing of transcripts encoding mechanotransduction proteins (Martino, Perestrelo et al., 2021). In contrast, glucocorticoid counteracts mechanotransduction in human skin fibroblasts cells by upregulating a lncRNA that promotes decay of mRNAs encoding mechanosensory proteins (Zhu, Li et al., 2021). Finally, the *filamin A* (*FLNA*) transcript edited by ADAR leads to a FLNA variant with increased actin crosslinking ability and thus increases cell stiffness, which could potentially alter cellular mechanotransduction (Jain, Weber et al., 2022). This collection of studies in addition to our present study of RNP1A revealing an intersection between mRNAs, CK machinery, and dynamic cell shape change processes like macropinocytosis that RNP1A provides strongly support the interconnectedness of these cellular sub-systems.

## Materials and Methods

### Expression plasmids

Generation of the plasmids for GFP–mCherry (linked), and GFP-and/or mCherry-tagged fusions of RNP1A, tubulin, Myosin II, Myosin II 3xAsp, Myosin II 3XAla, Cortexillin I, IQGAP1, IQGAP2 have been described previously (Ngo et al., 2016; Effler, Kee et al., 2006; Wheatley & Wang, 1996; Kee et al., 2012; Luo et al., 2013). GFP-DlpA was obtained from NBRP Nenkin. GFP-Atg18 was obtained from Dicty Stock Center (Otto et al., 2004).

### Cell culture and maintenance

*Dictyostelium discoideum* cells were maintained in Hans’ enriched HL-5 media (1.4X HL-5 media with 8% FM, penicillin and streptomycin) at 22°C in petri dishes (Fisher Scientific; FB0875712). A full list of strains used in this study can be found in **Appendix Table S3**.

Cells transformed with plasmids were grown in HL-5 media supplemented with corresponding selection drug (10 μg/ml or 15 μg/ml G418; 40 μg/ml hygromycin).

### Generation of *rnp*1A knockdown cell line

We first attempted a genetic deletion using CRISPR. For this, the CRISPR guide RNAs used were as follows:

Guide 1: 5’-CCAACAAAAATTCTGTGGGC-3’

Guide 2: 5’-ATTCTGTGGGCTGGAGTAGT-3’

For the *rnp*1A knockdown strains, we used the Ax3(Rep orf+; orfJ) wild type strain as the parental background. Knockdowns were generated using RNA hairpin targeting the 3’ of cDNA sequence. Using the pLD1A15SN plasmid, the following sequences were cloned into the plasmid to create the RNA hairpin:

Sense strand: 5’-CGGTTTCGTCGAATTCGATGATGTTGCCAATCAACAAAAAGGTCTCACCCTTAACAAACTCTCTGTTGAAAGTAGAGAACTCTCTGTTAAAATCGCTTTAGTTCCAGAACCAAGAGATGCCACC GCAACTACTCCAGATGTTACCACTACCGCT-3’

Anti-sense strand: 5’-AGCGGTAGTGGTAACATCTGGAGTAGTTGCGGTGGCATCTCTTGGTTCTGGAACTAAAGCGATTTTAACAGAGAGTTCTCTACTTTCAACAGAGAGTTTGTTAAGGGTGAGACCTTTTTGTTGATTGGCAACATCATCGAATTCGACGAAACCGAAACCTTTGCTTCTGTTGGTGTGTTTGTTGACAATGACATGAGCACTCTTTGGTGAGCAATCTTTGAAGGTTTCTAATAATTTAACATCATCAAAAGAGTATGGAATGTTTCTGACAAAGAGG GTAGTGGTACTTTGTTGTCTGTCAGCAGTGTTGGCAGCTGG-3’

Cells were transformed with 1 µg empty control plasmid or plasmid containing the *rnp1A* hairpin on day 0 and incubated in media without selection drug (G418) for two days. On day 2, the media was replaced to add 15 µg/mL G418. On day 4, the media was changed to reduce drug selection to 7.5 µg/mL G418. On day 6, media was changed again to reduce G418 to 5 µg/mL. From here on, media was changed with 5 µg/mL G418 every two days until cell colonies were visible on the substrate of cell culture dishes. Then, media was replaced every other day to increase G418 concentration by 1 µg/mL until 10 µg/mL, the concentration that was maintained from then on. *rnp*1A mRNA knockdown was verified and quantified by real time PCR and protein knockdown was verified and quantified by western blot.

The doxycycline-inducible *rnp1A* knockdown cell line was generated using doxycycline-inducible expression vector (pDM310), containing the same *rnp1A* hairpin sequence as above. 10 μg/mL doxycycline was used to induce expression and the induction lasted for either 48 hours or 100 hours. Knockdown was verified by qRT-PCR and western blotting.

### Growth assays

For suspension growth, cells were grown in suspension culture in flasks with a starting cell density of 10^4^ cells/mL. For the substrate growth assay, cells were grown in 8-well imaging chambers. Cell counts were taken every 24 hours. Cell growth rate were determined from the exponential growth phase. Relative growth rates were calculated as growth rate of individual samples divided by the average growth rates of control cell lines.

### Adhesion assays

*Dictyostelium* cells were plated on non-tissue culture 6-well plates for an hour at a 10^6^ cells/mL density. Then, 200 μL samples were taken from the culture media from each well, and cell counts were measured. The 6-well plates were shaken on a shaking platform at 50, 75, or 100 rpm for 30 min at room temperature. After shaking, 200 μL samples were again collected from the culture media from each well, and cell counts were measured. The fraction of cells non-adherent was calculated as the density of cells in culture media divided by the total density of cells on the plates. The change in fraction of cells that were non-adherent was calculated by subtracting the control fraction of non-adherent cells from mutant fraction of non-adherent cells.

### Immunofluorescence and imaging

Cells were plated in 8-well slide chamber (Lab-Tek) at a density of 1.5×10^6^ cells/mL for 30 min. Cell culture media was removed, and cells were fixed in -20°C acetone for 5 min or 2% PFA for 10 min. Cells were washed with PBST (1xPBS plus 0.05% Triton X-100) one time and then incubated with 3% BSA in PBST for 1 hour at room temperature. Primary antibody immunostaining was performed using a 1:10,000 dilution in 3% BSA in PBST at 4°C overnight. Then cells were washed three time with PBST. Secondary antibody immunostaining was performed using 1:3000 dilution in 3% BSA in PBST for one hour at room temperature, with the exception of the Rhodamine-phalloidin staining which was performed using 1:200 dilution 3% BSA in PBST for one hour at room temperature. Cells were again washed with PBST twice before Hoechst staining (10 μg/mL in 3% BSA in PBST) for 10 min at room temperature. Slides were then mounted with Invitrogen ProLong Diamond Antifade Mountant (P36961). Images were acquired with an Olympus IX71 epifluorescence microscope or with a Zeiss AxioObserver. Images were processed and quantified using ImageJ.

### Nuclei per cell measurement

Immunofluorescence staining was done as previously mentioned. Cells in exponential growth phase were grown in petri dishes prior to fixation. Nuclei per cell number was manually counted. Statistical analysis was performed using comparison of proportions.

### Cytokinesis imaging

Live cell images were taken at 2 sec intervals for 5-10 min. Time-lapse imaging was initiated at the beginning of cleavage furrow formation and terminated after cell division completed or failed. Cleavage furrow or intercellular bridge length, diameter, and the distance between two poles of the dividing cells were manually tracked and measured using ImageJ.

### Generation of RNP1A antibody and western blotting analysis

RNP1A antibody was developed by AbClonal. Full length RNP1A peptide was used as antigen. Four immunizations were performed on rabbits every 2 weeks. After the 4^th^ immunization, bleeds were collected from the rabbits, and polyclonal antibody was purified from the bleed.

For Western blotting analysis, cell lysates were prepared by boiling cells in SDS sample buffer, electrophoretically separated on SDS-polyacrylamide gels, and transferred to nitrocellulose membranes. Proteins were detected by each individual antibody. A list of antibodies used can be found in **Appendix Table S4**. Images were acquired on a LiCor Odyssey CLx system. Quantification of protein expression level was done by quantifying intensity of each band on the blot, background subtracted, and normalized against total protein amount measured from the Coomassie staining of a replica gel.

### Micropipette aspiration (MPA), effective cortical tension and mechanoresponsiveness quantification

MPA was performed with equipment set up as previously described (Kee and Robinson, 2013). Cells were seeded in an imaging chamber at around 10% confluency at least 30 minutes before imaging. The micropipette (∼5 μm diameter) was stabilized at the bottom of the cell chamber. To attach cells to the pipette tip, a small aspiration pressure was applied. Then, aspiration pressure was increased gradually to the equilibrium pressure ΔP, where the length of the cell inside the pipette (L_p_) is equal to the radius of the pipette (R_p_). The cell was then released from the pipette tip. After resting for a couple of minutes, a second measurement was performed on the same cell. Effective cortical tension is quantified using the Young-Laplace equation:

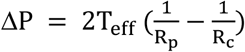

where

ΔP = aspiration pressure that produces cell deformation

T_eff_ = effective cortical tension

R_p_ = length of the cell inside the pipetter

R_c_ = pipette radius

Effective cortical tension for each cell is represented by the average from the two measurements.

To quantify mechanoresponse, cells were slowly aspirated to 0.80 nN/μm^2^ and held for at least 100 seconds. Mechanoresponsiveness was then calculated as the ratio of the background-corrected mean signal intensity of the cortex inside the pipette (I_p_) to that of the opposite cortex outside the pipette (I_o_).

### Fluorescence correlation spectroscopy and cross-correlation spectroscopy

Fluorescence Correlation Spectroscopy (FCS) and Fluorescence Cross-Correlation Spectroscopy (FCCS) were performed as previously described (Kothari et al., 2019). Specifically, cells expressing the corresponding fluorophore-labeled proteins were plated on imaging chambers at least 30 minutes before imaging. We used co-expressed soluble GFP and mCherry as negative control, and expressed GFP attached to mCherry by a 5 amino-acid flexible linker as positive control. System calibration was performed using 100 nM Rhodamine. Experiments were performed on a Zeiss AxioObserver with 780-Quasar confocal module and FCS capability using a C-Apochromat 40× water objective. The “apparent *in vivo*” *K*_*D*_ was calculated using the following equation:

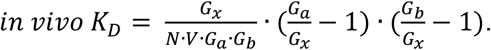

where G_x_ = cross-correlation of two fluorophores

G_a_ = auto-correlation for mCherry

G_b_ = auto-correlation for GFP

N = Avogadro’s number

*V* = confocal volume

Image data was processed with Zen (black) software using the FCCS analysis module.

### Agarose overlay and microtubule length quantification

Agarose overlay was performed as described previously (Kee et al., 2012). Specifically, thin sheets of 2% agarose gel in water were prepared as follows. Two 22 mm x 22 mm cover glasses (Fisherbrand 12542B) were placed at each end of a 22 mm x 60 mm microscope cover glass (Fisherbrand 12545J). 1 mL of dissolved agarose solution was placed onto the center of the 22 mm x 60 mm microscope cover glass, and a second 22 mm x 60 mm microscope cover glass was immediately placed on top to spread the agar evenly between two ends of the cover glass. This method produces thin agarose sheets of around 0.15 mm thick. Agarose sheets were cut into smaller pieces to fit into imaging chambers (Lab-Tek 155409). Prior to imaging, cells were plated onto imaging chambers and allowed to sit for 30 min. Cell culture media was removed, and pieces of 2% agarose sheets were placed directly on top of cells in the imaging chamber.

To disassemble microtubules, cells were treated with 8 μM thiabendazole for 30 min and a negative control was run in parallel with the same volume of DMSO (vehicle). The GFP-labeled microtubules were traced manually and lengths measured using ImageJ.

### Total Internal Reflection Fluorescence Imaging

Total Internal Reflection Fluorescence (TIRF) microscopy was performed on an Olympus IX81 microscope. Movies were acquired using a 60X objective and a 1.6X optivar. TIRF movies of GFP-tubulin were captured at 1 sec intervals for a duration of 30 sec. The number of GFP-tubulin cortical contacts were manually traced and quantified in ImageJ for each cell at every second over the span of a 30 sec movie.

### RNA-seq

RNA-seq was performed in collaboration with Novogene. Specifically, two replicates of total RNA was harvested from each of the two independent rounds of RNA hairpin knockdown of *rnp1A* (both wild type control and *rnp1A* hairpin cell lines; the two independent rounds of RNA hairpin knockdowns are presented as R1 and R2). A total amount of 1 μg RNA per sample was used as input material for the RNA sample preparations. Sequencing libraries were generated using NEBNext® UltraTM RNA Library Prep Kit for Illumina® (NEB, USA) following the manufacturer’s recommendations, and index codes were added to attribute sequences to each sample. Briefly, mRNA was purified from total RNA using poly-T oligo-attached magnetic beads. Fragmentation was carried out using divalent cations under elevated temperature in NEBNext First Strand Synthesis Reaction Buffer (5X). First strand cDNA was synthesized using random hexamer primer and M-MuLV Reverse Transcriptase (RNase H-). Second strand cDNA synthesis was subsequently performed using DNA Polymerase I and RNase H. Remaining overhangs were converted into blunt ends via exonuclease/polymerase activities. After adenylation of the 3’ ends of DNA fragments, NEBNext Adaptor with a hairpin loop structure were ligated to prepare for hybridization. To select cDNA fragments of preferentially 150∼200 bp in length, the library fragments were purified with AMPure XP system (Beckman Coulter, Beverly, USA). Then 3 μl USER Enzyme (NEB, USA) was used with size-selected, adaptor-ligated cDNA at 37 °C for 15 min followed by 5 min at 95 °C before PCR. Then PCR was performed with Phusion High-Fidelity DNA polymerase, Universal PCR primers and Index (X) Primer. At last, PCR products were purified (AMPure XP system) and library quality was assessed on the Agilent Bioanalyzer 2100 system. The clustering of the index-coded samples was performed on a cBot Cluster Generation System using PE Cluster Kit cBot-HS (Illumina) according to the manufacturer’s instructions. After cluster generation, the library preparations were sequenced on NovaSeq 6000 (PE150) and paired-end reads were generated. Raw data (raw reads) in FASTQ format were firstly processed through fastp. Paired-end clean reads were mapped to the reference genome using HISAT2 software. Featurecounts was used to count the read numbers mapped to each gene. And then the RPKM of each gene was calculated based on the length of the gene and the read count that was mapped to this gene. Differential expression analysis was performed using DESeq2 R package. Gene Ontology (GO) enrichment analysis of differentially expressed genes was implemented by the clusterProfiler R package. We used clusterProfiler R package to test the statistical enrichment of differential expression genes in KEGG pathways.

### RNA extraction and qRT-PCR

RNA extraction was done with TRIzol™ reagent (Thermo Fisher 15596026), following the manufacturer’s instruction.

qRT-PCR was performed using Verso 1-step RT-qPCR kit (AB4104A). The primers used for qRT-PCR are listed in **Appendix Table S5**. All experiments were conducted on a BIO-RAD CFX Opus 96 Real-Time PCR system. The program used was the following: Reverse Transcription - 50°C for 15 minutes, 95°C for 15 minutes; PCR - 95°C for 15 seconds, 60°C for 30 seconds, 72°C for 30 seconds; Melt curve generation - 95°C for 30 seconds, 60°C for 30 seconds, and then increased by 0.5°C per 10 seconds. Quantification of gene expression from qRT-PCR was performed by calculating the differences in C_t_ value between target genes and control gene (*abcF4*; *abcF4* did not exhibit gene expression changes from RNA-seq) for each cell line, and then normalized against wild type control cell line.

### CLIP-seq

CLIP-seq protocol was adapted from a previous publication (Muller, Windhof et al., 2013). Specifically, cells were first washed one time with phosphate buffer and then plated in petri dishes for UV crosslinking at 250 mJ/cm^2^. Cells were resuspended in modified RIPA buffer (50 mM Tris-HCl pH 7.5, 150 mM NaCl, 0.5% NP40, 0.5% sodium deoxycholate, 1 mM EDTA), and then sonicated at 30% amplitude for 4 minutes (30 seconds on, 30 seconds off). Samples were then centrifuged for three time at 20,000xg for 15 minutes at 4°C, and the supernatant was removed. Supernatant was pre-incubated with 250 µL Sephadex G-50 beads in TE buffer. 40 µL was collected as input one. And then supernatant was incubated with 50 µL GFP-trap beads (Chromotek, gta) overnight. On the next day, beads were washed once with RIPA stringency A buffer (50 mM Tris-HCl pH 7.5, 1 M NaCl, 1% NP40, 1% sodium deoxycholate, 1 mM EDTA, 0.1% SDS, 2 M Urea), twice with RIPA stringency B buffer (50 mM Tris-HCl pH7.5, 1 M NaCl, 1% NP40, 1% sodium deoxycholate, 1 mM EDTA, 0.1% SDS, 1 M Urea). 40 µL was collected as input two. Beads were then washed once with equilibration buffer (50 mM Tris HCl pH 7.5, 300 mM NaCl, 0.5% NP40, 0.5% sodium deoxycholate, 1 mM EDTA, 0.1% SDS). Finally, beads were added to elution buffer (50 mM Tris-HCl pH 7.5, 300 mM NaCl, 0.5% NP-40, 0.5% sodium deoxycholate, 5 mM EDTA, 0.1% SDS, 1 mg/mL proteinase K) and incubated at 42°C for one hour followed by 56°C for four hours. RNA was extracted with TRIzol™ reagent (Theormo Fisher 15596026). 3 replicates per cell line (control: wild type cells expressing GFP; sample: wild type cells expressing RNP1A-GFP) were used in downstream sequencing and analysis. RNA samples were converted to double stranded cDNA using the Ovation RNA-Seq System v2.0 kit (Tecan, Männedorf, Switzerland), which utilizes a proprietary strand displacement technology for linear amplification of mRNA without rRNA/tRNA depletion as per the manufacturer’s recommendations. This approach does not retain strand specific information. Quality and quantity of the resulting cDNA was monitored using the Bioanalyzer High Sensitivity kit (Agilent) which yielded a characteristic smear of cDNA molecules ranging in size from 500 to 2000 nucleotides in length. After shearing 500 nanograms of cDNA to an average size of 250 nucleotides with the Covaris S4 (Covaris Inc., Woburn, MA) library construction was completed with the Truseq Nano kit (Illumina; San Diego, CA) according to the manufacturer’s instructions. mRNA libraries were sequenced on an Illumina Novaseq 6000 instrument using 150bp paired-end dual indexed reads and 1% of PhiX control. Reads were aligned to the *Dictyostelium discoideum* reference genome dicty2.7.51. rsem-1.3.0 was used for alignment as well as generating gene expression levels. The ‘rsem-calculate-expression’ module was used with the following options: --star, --calc-ci, --star-output-genome-bam, --forward-prob 0.5. Differential expression analysis and statistical testing were performed using DESeq2 software. Transcripts identified to bind to RNP1A meet the threshold of padj ≪ 0.1.

### Macropinocytosis measurements

Cells were seeded onto 8-well imaging chambers (Nunc Lab-Tek) at a density of 5×10^5^ cells/mL for 30 minutes before imaging. Imaging was performed on a Zeiss LSM 780 FCS confocal microscope. Prior to imaging, TRITC-Dextran (Millipore-Sigma; 65-85 kDa) was added to cells at a final concentration of 1 mg/mL. Images were acquired every 30 sec for a total duration of 10 or 20 min. We randomly chose 10-20 cells for analysis. Mean TRITC signal intensity at certain time points during the movie for an individual cell was measured, corrected for background, and then normalized to that of the first frame. In the end, mean and standard errors of normalized TRITC intensity over time were calculated for each individual cell line for comparison. Starvation was induced by keeping cells in the developmental buffer (DB) for 2 hours.

To measure macropinocytotic crown areas, images were taken every second after adding 1 mg/mL TRITC-Dextran for a total duration of 2 min. Five to ten cells were randomly chosen from each movie for measurement. We identified the first frame where the macropinocytotic crown membrane closed, and the crown at the identified frame was manually traced to measure the area. The number of macropinocytotic events per cell over the span of 2 min was also recorded for comparison between cell lines. For visualization of RNP1A, DlpA and Contractility Kit (CK) proteins localization during macropinocytosis, GFP or mCherry tagged proteins were expressed in different cell line backgrounds, and movies of macropinocytosis were taken as described above.

To measure the rate of loss of internalized TRITC-Dextran in cells, cells were seeded and TRITC-Dextran was added as described above. After around 10 min, cells were washed with HL5 media for three times and then fresh HL5 media was added to the chamber to remove the remaining TRITC-Dextran in the media. Images were acquired every 30 sec for a total duration of 60 min. 20 cells were randomly chosen for analysis described as above.

### Statistical analysis

All statistical analysis was performed with Kruskal–Wallis followed by Wilcoxon-Mann–Whitney test, unless otherwise specified. Annotation used in figures: *,P≤0.05; **,P≤0.01; ***,P≤0.001; ****,P≤0.0001; n.s., not significant.

## Acknowledgements

We thank AbClonal for producing the RNP1A antibody, Novogen for performing RNA-seq experiment and analysis, Johns Hopkins Experimental and Computational Genomics Core for performing CLIP-seq and analysis, and the Johns Hopkins School of Medicine Microscope Core Facility to provide instructions for and access to microscopes and analysis software.

We thank NBRP Nenkin for providing *dlpA* null cells and GFP-DlpA plasmids, Dicty Stock Center for GFP-Atg18 plasmid, and Developmental Studies Hybridoma Bank for providing various antibodies for this study.

Lastly, we thank the entire Robinson lab, especially Xuerui Wang, for their inputs and suggestions.

This work was supported by NIH grant GM66817.

## Data availability

The datasets produced in this study are available in **Datasets EV1** and **EV2**, as well as in the following database:

- RNA-seq data: Gene expression Omnibus GSE207131(https://www.ncbi.nlm.nih.gov/geo/query/acc.cgi?acc=GSE207131).

## Author contributions

Y.L..: Conceptualization; Data curation; Formal analysis; Investigation; Methodology; Validation; Visualization; Writing-original draft; Writing-review&editing. J.L.: Conceptualization; Data curation; Formal analysis; Investigation. L.N.: Data curation; Formal analysis; Methodology. L.N. conducted cortical tension and mechanoresponsiveness measurements as well as corresponding data analysis. D.N.R.: Conceptualization, Funding acquisition, Resources, Supervision, Writing-review&editing.

## Expanded View Figure legends

**Expanded View Figure 1. F-actin amount is reduced in *rnp1A* knockdown and increased in RNP1A overexpressing cells**. (A) Fluorescence images of F-actin in wild type control, *rnp1A* knockdown and RNP1A overexpressing cells labeled with Rhodamine-phalloidin. Cells were fixed with 2% PFA. Scale bar, 10 µm. (B) Quantification of Rhodamine intensity per cell in wild type control, *rnp1A* knockdown and RNP1A overexpressing cells. Mean Rhodamine intensity of each cell is measured and normalized against the average of that in wild type control cells. Data were pooled from 20 cells per cell line. Statistical analysis was performed with Kruskal–Wallis followed by Wilcoxon-Mann–Whitney test. **,P≤0.01;***,P≤0.001.

**Expanded View Figure 2. RNA-seq results from *rnp1A* knockdown cells (replicate 2)**. (A) Volcano plot of differentially expressed genes from RNA-seq on *rnp1A* knockdown (replicate 2) is shown. Differentially expressed genes are at least two-fold up-regulated or down-regulated, and their corresponding padjs are equal or smaller than 0.1. Most significantly up- or down-regulated genes are labeled with DDB_ID or gene name. (B) Gene ontology analysis of up-regulated genes from RNA-seq of *rnp1A* knockdown cells (replicate 2). (C) Gene ontology analysis of down-regulated genes from RNA-seq of *rnp1A* knockdown cells (replicate 2).

**Expanded View Figure 3. GFP-Atg18 aggregation is reduced in *rnp1A* doxycycline-inducible knockdown cells**. (A) Images show GFP-Atg18 localization in control and *rnp1A* doxycycline-inducible knockdown cells under 2% agarose gel compression at 0, 2.5, and 4.5 hours. White arrowheads point to example GFP-Atg18 aggregates. Scale bar, 10 µm. (B) Quantification of number of GFP-Atg18 aggregates in control and *rnp1A* doxycycline-inducible knockdown cells under 2% agarose gel compression at 0, 2.5, and 4.5 hours. Number of GFP-Atg18 aggregates are respectively normalized against cell area or cell number. Data were pooled from measurements from 60 cells per cell line. Error bars indicate standard errors.

**Expanded View Figure 4. Over- and under-assembly of Myosin II reduced TRITC-Dextran uptake**. (A) TRITC-Dextran uptake in *myosin II* null cells expressing GFP, GFP-Myosin II, GFP-3xAla, GFP-3XAsp, and RNP1A-GFP. TRITC-Dextran intensity was quantified by mean TRITC intensity of each cell, background subtracted, normalized to cell area, and then normalized to that at first time point. Scale bar, 10 µm. Data were pooled from measurements from the numbers of cells indicated in the figure. Error bars indicate standard errors. (B) Quantification of macropinocytotic crown area and number of macropinocytotic events per cell over the span of 2 minutes. Macropinocytotic crowns were manually traced as shown in yellow circles in images. Scale bar, 5 µm. Data were pooled from the number of cells indicated in the figure. Statistical analysis was performed with Kruskal–Wallis followed by Wilcoxon-Mann–Whitney test. *, P≤0.05.

